# Quantitative and qualitative evaluation of the impact of the G2 enhancer, bead sizes and lysing tubes on the bacterial community composition during DNA extraction from recalcitrant soil core samples based on community sequencing and qPCR

**DOI:** 10.1101/365395

**Authors:** Alex Gobbi, Rui G. Santini, Elisa Filippi, Lea Ellegaard- Jensen, Carsten S. Jacobsen, Lars H. Hansen

## Abstract

Soil DNA extraction encounters numerous challenges that can affect both yield and purity of the recovered DNA. Clay particles lead to reduced DNA extraction efficiency, and PCR inhibitors from the soil matrix can negatively affect downstream analyses when applying DNA sequencing. Further, these effects impede molecular analysis of bacterial community compositions in lower biomass samples, as often observed in deeper soil layers. Many studies avoid these complications by using indirect DNA extraction with prior separation of the cells from the matrix, but such methods introduce other biases that influence the resulting microbial community composition.

To address these issues, a direct DNA extraction method was applied in combination with the use of a commercial product, the G2 DNA/RNA Enhancer^®^, marketed as being capable of improving the amount of DNA recovered after the lysis step. The results showed that application of G2 increased DNA yields from the studied clayey soils from layers between 1.00 and 2.20 m below ground level.

Importantly, the use of G2 did not introduce bias, as it did not result in any significant differences in the biodiversity of the bacterial community measured in terms of alpha and beta diversity and taxonomical composition.

Finally, this study considered a set of customised lysing tubes for evaluating possible influences on the DNA yield. Tubes customization included different bead sizes and amounts, along with lysing tubes coming from two suppliers. Results showed that the lysing tubes with mixed beads allowed greater DNA recovery compared to the use of either 0.1 or 1.4 mm beads, irrespective of the tube supplier.

These outcomes may help to improve commercial products in DNA/RNA extraction kits, besides raising awareness about the optimal choice of additives, offering opportunities for acquiring a better understanding of topics such as vertical microbial characterisation and environmental DNA recovery in low biomass samples.

## Introduction

The complex chemical and physical structure of soil greatly influences the binding strength of DNA to its particles. Soil factors, such as clay type and content, concentration and valence of cations, the amount of humic substances and pH, dictate the adsorption of DNA into the soil matrix [1, 2]. Clay minerals effectively bind DNA and other charged molecules, such as Ca^2+^, Mg^2+^ and Al^3+^. This binding effect can be either direct, due to the positively charged edges of the clay particles, or indirect due to the neutralising bridge between negative charges on clay surfaces and DNA macromolecules caused by the presence of polyvalent cations (Ca^2+^, Mg^2+^ and Al^3+^) [2-4]. The increased binding of DNA to soil particles significantly reduces degradation of this biological material by extracellular microbial DNases and nucleases [5-7]. However, the same binding force that protects DNA from degradation reduces the amount of DNA recovered during DNA extraction. Appropriate DNA recovery from the soil matrix is also affected when the soil layers, especially those located below the topsoil, contain lower biomass and hence smaller DNA quantities. Indeed, deeper soil layers generally have a lower carbon content, lower nutrient concentrations and consequently lower microbial biomass [8, 9]. Despite the decrease in microbial biomass with soil depth, the hitherto poorly characterised communities dwelling in the deeper layers of the soil perform important roles in carbon sequestration [10], nutrient cycling [11, 12], mineral weathering and soil formation [13, 14], contaminant degradation [13] and groundwater quality [11].

Depending on the methodological approach applied, DNA extraction methods can be divided into direct and indirect methods. In direct extraction protocols, lysis is the first step and the microorganisms are treated with the matrix. During indirect extraction, however, the first step involves the detachment of the microbes from the soil matrix, generally conducted in a liquid media or supplemented by the use of density centrifugations such as in Nycodenz extractions [15]. Both these methods introduce a different extraction bias [16]. During direct extractions from inorganic soils that are particularly rich in clay, the DNA liberated from the microbial cells is quickly absorbed onto clay particles, preventing complete recovery. During indirect extractions, however, the bias is due to the different efficacy of the separation treatment on specific microbes, which may enrich one particular microbial fraction over another [17]. These complications may impede the extractions from soil and sediment layers, particularly from lower depths, and thus hamper the study of topics such as vertical microbial characterisation and environmental ancient DNA. The importance of obtaining high DNA yields is also related to greater representativeness of the soil gene pool, reducing the bias introduced into the successive analyses [18, 19].

Although many authors working with soil or sediments have opted for indirect DNA extraction methods to overcome these problems [20-23], the present study focused on a direct approach. Direct extraction methods have been shown to recover the greatest diversity in terms of the number of OTUs, especially if based on mechanical lysis such as bead beating when compared with other lysis methods [24]. With the aim of reducing the retention capacity of clay particles, the commercial product G2 DNA/RNA Enhancer^®^ (Ampliqon A/S, Odense, Denmark), hereafter referred to as G2, has been introduced into lysing matrix tubes. The G2 product is marketed as being able to improve the amount of DNA recovered after the lysis step [25]. G2 is a product made from freeze-dried highly mutagenised salmon sperm DNA [25] that adsorbs, like environmental DNA, to the clay particle before cell lysis [26]. However, unlike salmon sperm DNA, the G2 enhancer is not amplifiable in downstream polymerase chain reaction (PCR) applications [25].

The primary objective of the present study was to test the effects of the commercial product G2 on DNA yield and the diversity of the bacterial community from silty clay soil samples from layers between 1.00 and 2.20 metres below ground level (mbgl), thus aiming to develop an improved and bias-reduced direct DNA extraction protocol for this type of challenging sample. In addition to the effect of G2 on DNA yield, tests were conducted using both customised and commercially available DNA extraction kits. Customised tubes for evaluating possible influences on the DNA yield were prepared using different bead sizes and amounts, along with different plastic lysing tubes from different suppliers.

## Material and methods

### Soil core sampling

Soil sampling was performed in September 2016 at a vineyard located in the municipality of La Horra in the Ribera del Duero region in Spain (41°43’46.82"N, 3°53’29.85"W). The sampling approach was designed to obtain undisturbed soil cores, allowing later sub-sampling for DNA analysis at specific soil core depths. To do so, a Fraste Multidrill model PL was applied for intact soil core sampling using a hydraulic hammer. This model is typically used for standard penetration test (SPT) analysis, but in this case it was fitted with a PVC tube adapted internally to the metal probe rod to allow undisturbed soil core recovery. The soil cores were recovered in the PVC tubes with a 2.5” diameter in lengths of 0.60 m. The cores were then sealed at both ends, labelled and stored in a cold room (4-6°C) until further sub-sampling.

### Soil sub-sampling and homogenisation

Sub-sampling was performed by opening the cores and recovering soil from specific depths and putting the samples into sterile 15 ml plastic tubes. Sub-samples were taken from the central, untouched part of the cores collected using a sterile spatula and tweezers. After sub-sampling, the 15 mL plastic tubes were promptly frozen at −18 °C, shipped to Denmark and kept frozen until DNA extraction.

To obtain a homogenous soil sample for distribution between the DNA extraction tubes, a composite soil sample was produced by mixing 22.5 g of soil. This pool was composed by mixing 1.5 g of soil from 15 selected soil sub-samples representing soil depths ranging from 1.00 to 2.20 mbgl, a depth range chosen on the basis of previous pilot studies. The pilot studies showed that below the depth of approximately 2.20 m, no measurable DNA could be recovered without using G2, which would make a comparison of microbial communities between the extractions with and without G2 unviable. To proceed with the DNA extraction, 0.4 g of the homogenised soil pool was placed in 51 lysing tubes. The DNA extraction followed the protocol of the FastDNA^®^ Spin Kit for Soil (MP Biomedicals, LLC, Solon, OH, USA), but was modified with regard to the lysing tubes, as summarised in Table 1.

**Table 1.** All the lysing tubes used in the experiments.

| **Series** | ***n*** | ***Tube*** | ***Beads*** | ***G2*** |
| --- | --- | --- | --- | --- |
| a* | 5 | MP-Bio | Mixed | No |
| b | 5 | MP-Bio | Mixed | Yes |
| c | 5 | MP-Bio | 1.4 mm | No |
| d | 5 | MP-Bio | 0.1 mm | No |
| e* | 5 | Ampliqon | 1.4 mm | Yes |
| f* | 5 | Ampliqon | 0.1 mm | Yes |
| g | 5 | Ampliqon | 1.4 mm | No |
| h | 5 | Ampliqon | 0.1 mm | No |
| i | 5 | Ampliqon | Mixed | No |
| A-NEG+G2 | 3 | MP-Bio | Mixed | Yes |
| A-NEG | 3 | MP-Bio | Mixed | No |

\*Products already commercialised, while the remainder are customised preparations. A-NEG+G2 represents a negative control of the extraction with G2 added; A-NEG represents a negative control of the extraction using the kit's lysing tube without a sample. With mixed beads, the intention was to have an exact proportion of 0.1 and 1.4 mm beads plus a large 4 mm glass bead. When mixed beads are used, the final weight of the beads is greater than the single type beads. It was decided not to modify this parameter since the test was intended to compare already existing commercial products. The customised preparations were made to evaluate the statistical effect of the different variables involved.

### DNA quantification qPCR, library preparation and sequencing

For this study, different types of lysing tubes were prepared using commercial products made by MP Biomedicals and Ampliqon in order to test the effect of plastics, beads and G2. All the preparations are summarised in Table 1.

Following the extractions, the DNA yields were measured using Qubit^®^2.0 fluorometer (Thermo Scientific™). Qubit measurements were taken from all the tube preparations in triplicates and the average results reported in Table in S1 Supporting Information. All PCR reactions were prepared using UV sterilised equipment and negative controls were run alongside the samples.

The qPCR with primers targeting the 16S rRNA gene was carried out on a CFX Connect™ Real-Time PCR Detection System (Bio-Rad). The primers used were 341F (TCGTCGGCAGCGTCAGATGTGTATAAGAGACAG-CCTAYGGGRBGCASCAG) and 806R (GTCTCGTGGGCTCGGAGATGTGTATAAGAGACAG-GGACTACNNGGGTATCTAAT) [27], complete with adapters for Illumina MiSeq sequencing.

Single qPCR reactions contained 4 µL of 5x HOT FIREPol^®^ EvaGreen^®^ qPCR Supermix (Solis BioDyne, Tartu, Estonia), 0.4 µL of forward and reverse primers (10 µM), 2 µL of bovine serum albumin (BSA) to a final concentration of 0.1 mg/mL, 12.2 µL of PCR grade sterile water and 1 µL of template DNA. A standard curve consisting of dilution series of 16S standard was prepared from DNA extracts of *Escherichia coli* K-12, with seven 16S rRNA gene copies per genome [28]. The quantity of 10^-^1 16S standard was 8.45 × 10^7^ 16S genes/µL. Quantification parameters showed an efficiency of E=87.7 % and R^2^ of 0.997. The qPCR cycling conditions included initial denaturation at 95 °C for 12 min, followed by 40 cycles of denaturation at 95 °C for 15 sec, annealing at 56 °C for 30 sec, and an extension at 72 °C for 30 sec, with a final extension performed at 72 °C for 3 min.

Amplicon library preparation was performed by a two-step PCR, as described by Feld, Nielsen (29) and Albers, Ellegaard-Jensen (30) with slight modifications. Sample concentration was approximately 5 ng of DNA, and both PCRs were carried out using a Veriti Thermal Cycler (Applied Biosystems). In each reaction of the first PCR, the mix contained 12 µL of AccuPrime™ SuperMix II (Thermo Scientific™), 0.5 µL of forward and reverse primer from a 10 µM stock, 0.5 µL of bovine serum albumin (BSA) to a final concentration of 0.025 mg/mL, 1.5 µL of sterile water and 5 µL of template. The reaction mixture was pre-incubated at 95 °C for 2 min, followed by 33 cycles of 95 °C for 15 sec, 55 °C for 15 sec, 68 °C for 40 sec, with a final extension performed at 68 °C for 4 min. Samples were subsequently indexed by a second PCR using the following PCR protocol. Amplification was performed in 28 µL reactions with 12 µL of AccuPrime™ SuperMix II (Thermo Scientific™), 2 µL of primers complete with indexes and P7/P5 ends, 7 µL of sterile water and 5 µL of PCR1 product. The cycling conditions included initial denaturation at 98 °C for 1 min, followed by 13 cycles of denaturation at 98 °C for 10 sec, annealing at 55 °C for 20 sec, and extension at 68 °C for 40 sec, with a final extension performed at 68 °C for 5 min. The primer dimers formed in the PCR were removed, along with PCR components, using a clean-up step. In this step, HighPrep™ PCR reagent (MAGBIO) was used to selectively bind the DNA fragments to sequence according to the manufacturer’s protocol. PCR products were finally checked by electrophoresis on a 1.5 % agarose gel. Samples were then pooled in an equimolar amount of 10 ng and sequenced on Illumina MiSeq instrument using 2×250 paired-end reads with V2 Chemistry.

## Bioinformatics

Sequencing data were analysed and visualised using QIIME 2 v. 2017.9 [31]. Demultiplexed reads from the Illumina MiSeq were quality filtered using the plugin quality filter with default parameters of QIIME2 [32]. Reads were then denoised, chimera checked and dereplicated using a DADA2 denoise-paired plugin [33]. The output was rarefied to the lowest sample at 16047 reads using qiime feature-table rarefy [34]. Thereafter a multiple-sequence alignment was performed using MAFFT [35] and subsequently a phylogenetic tree generated using FastTree [36]. Alpha and beta diversity analyses were performed through a q2-diversity plugin [37] with the core-metrics-phylogenetic method on the rarefied sequence-variant table. For the alpha diversity, two different parameters were measured: richness and evenness. Richness was measured based on Faith-pd [38], while evenness [39] was reported through the Pielou score [40]. This produced box plots and PCoA plots visualised through Emperor [41]. Taxonomic assignments were performed using qiime feature-classifier classify-sklearn in which a pre-trained Naïve-Bayes classifier with Greengenes v_13.8 [42] was used. Taxa bar plots were built using the plugin qiime taxa bar plot with different filtered, unfiltered and grouped rarefied tables.

## Statistics

A statistical evaluation of the results was performed separately for DNA quantification and sequencing dataset. For DNA quantification, assessments based on one-way ANOVA followed by Scheffe’s test were performed. The ANOVA test indicates whether there is at least one statistically significant (p-value below 0.05) difference in the whole dataset. In contrast, Scheffe’s method indicates which group comparison is statistically significant (comparison score > critical Scheffe’s score). The critical value of Scheffe’s method is calculated starting from the F-critic of ANOVA test multiplied by (N-1), where N is the number of comparisons performed with Scheffe’s method. Scheffe’s test was used here because it is less sensitive to an unequal number of samples representing the different variables analysed. Furthermore Scheffe’s methods have been used to reduce false positive results due to type I errors when multiple comparisons are performed on the same dataset. Both tests were applied to the dataset of the DNA quantification obtained using Qubit and qPCR for the gene copy number. Since both results were consistent and G2 is mainly used to obtain DNA for PCR-based downstream applications, only the results of the qPCR are discussed. All statistical evaluations and visualisations regarding DNA amount and gene copy number quantification were performed in Microsoft Office Excel 2010.

Sequencing data after QIIME 2 pipeline processing (2.4) were statistically evaluated using the Kruskal-Wallis test for alpha and beta diversity, a non-parametric method substitute of ANOVA when the normal distribution of data cannot be assumed. The resulting p-value of the alpha diversity comparison was based on the medians of different parameters (richness and evenness) calculated between the different series analysed. Beta diversity analyses were performed using both Kruskal Wallis and PERMANOVA with 999 permutations. Finally a statistical evaluation was performed of differentially abundant features based on an analysis of composition of microbiomes (ANCOM). ANCOM computes Aitchinson’s [43] log-ratio of relative abundance for each taxon, controlling the false discovery rate (FDR) using the Benjamini-Hochberg procedure. This test is based on the assumption that few features change in a statistical way between the samples, and hence it is very conservative [44]. All these statistical tests were applied using QIIME2 v2017.9.

## Results

The main aim of the present study was to evaluate the impact of the commercial product G2 on DNA extraction from a silty clay soil layer between 1.00 and 2.20 mbgl. G2, freeze-dried inside the lysing tubes, was applied with the purpose of preventing or reducing the adsorption of environmental DNA onto the clay particles in the soil, thus improving the recovery of nucleic acids during cellular lysis. This paper presents the results of the impact of G2 on final DNA extraction efficiency and its possible impact on the composition of soil microbial communities. Furthermore the G2 effect was compared with other variables such as bead size and tube supplier in different commercialised and customised lysing tubes, as summarised in Table 1.

### G2 enhanced DNA recovery from deep soil layers

In order to test the influence of the presence of G2 on DNA yield, a series of extractions were set up using different bead size and tube combinations. Following DNA quantification *via* Qubit, the results were analysed. As seen in Fig 1A, the G2 component consistently improved the DNA yield (Scheffe’s score > 27.612 – Table in S2 Supporting Information). Fig 1A shows that there were differences between the different tube preparations, irrespective of the presence of G2. Different bead combinations and tubes without the addition of G2 were therefore tested. These variations could potentially be attributed to the different beads and the plastic of the tubes (Fig 1B). The mixed beads allowed greater DNA recovery compared to the use of either 1.4 mm or 0.1 mm beads, regardless of the type of plastic tubes used (Scheffe’s score > 27.612 – Table in S2 Supporting Information). The differences in the DNA yield between the 1.4 mm and 0.1 mm beads were less pronounced, including for the differences due to the plastic composition of the tubes. Mixed beads always allowed recovery of the highest DNA yield in all the tube preparations in which they were used.

**Fig 1.**
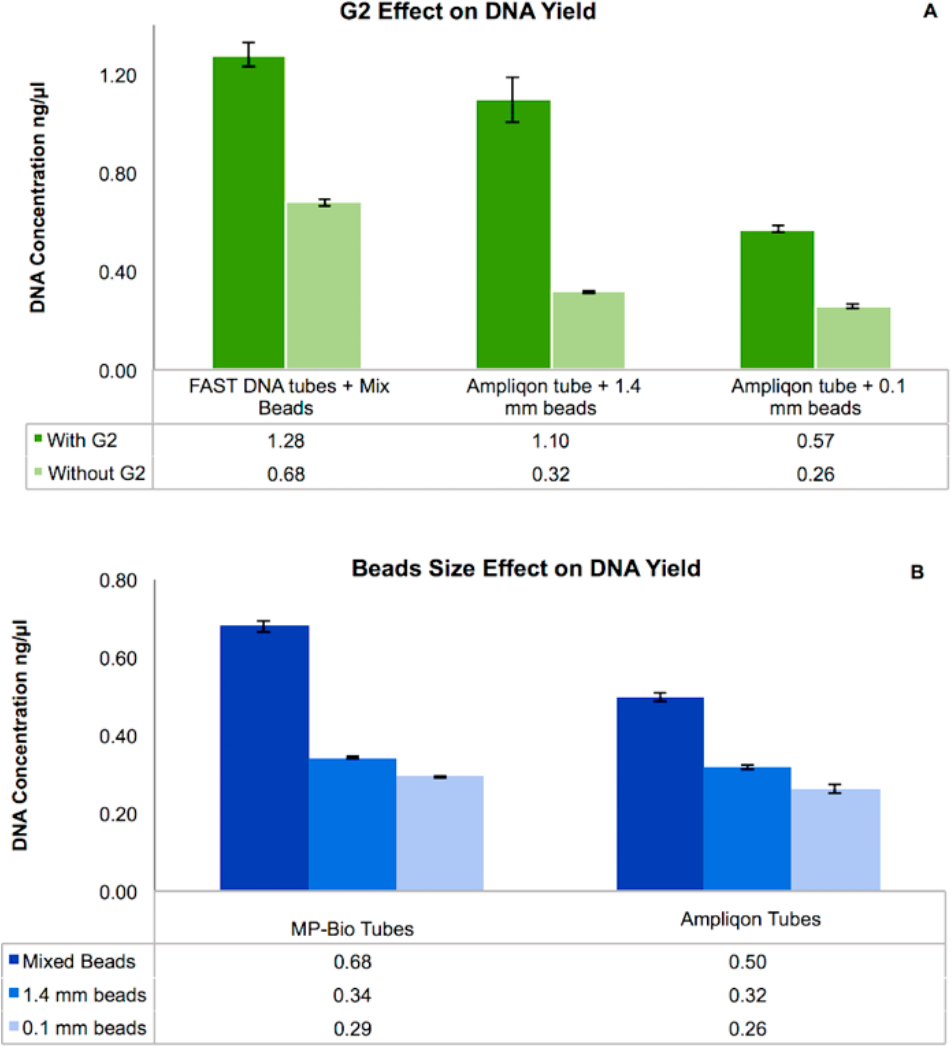
Results based on Qubit quantification after DNA extraction expressed as ng/µl. Comparison of three commercial products with and without G2. (B) Comparison of three different bead sizes and the two tube types, without added G2. Values shown are averages of n=3 independent measurements on n=5 biological replicates. Error bars are calculated from the standard deviation for each series. There was a statistical significance (Scheffe’s score > 27.612) for the three comparisons with and without G2 in test A, while for test B, it occurred in the comparisons between mixed X 1.4 mm beads and mixed X 0.1 mm beads, but not between 1.4 mm X 0.1 mm beads or between the two tube suppliers (Table in S2 Supporting Information).

### Greater DNA yield corresponded to a higher number of 16S genes

Considering that G2 is a DNA-based product, this could potentially affect the Qubit measurements since the signal recorded using a fluorescent dye is emitted when it binds dsDNA. In such a situation there might be an overestimation of soil DNA from the Qubit result. To examine whether this was the case, the copy number variation of a genetic marker, such as the 16S rRNA gene for bacteria, was tested via qPCR.

The results of the qPCR (Fig 2 and in Table in S3 Supporting Information) confirmed the trend observed with the DNA yield provided by the Qubit analysis. The ANOVA based on the qPCR results described the dataset with a p-value of 1.0147*E^-^36 (Table in S4 Supporting Information). This meant that in all the data produced by qPCR, taking into account the variance within and between groups, there was at least one difference that was statistically significant. However, since an ANOVA test does not identify where this difference is, Scheffe’s method was applied to compare the different combinations of G2, bead size and tube plastic (Table in S5 Supporting Information). Based on these results, it was evident that G2 always allowed the recovery of the highest number of genes, while mixed beads also improved gene recovery (Scheffe’s score > 26.117 – Table in S5 “a” Supporting Information). These results were also evident and consistent when looking at the DNA yield measured *via* Qubit, as reported in Table in S1 Supporting Information. The comparison of different plastic tubes (Table in S5 “e” Supporting Information) showed a non-significant difference. Instead, the beads used in lysis were shown to have a significant effect on DNA yield (Table in S5 “b” Supporting Information) and, as discussed below, also on microbial composition. From the Greater DNA yield corresponded to a higher number of 16S genes Scheffe’s statistics, it was possible to observe that the mixed beads from the original lysing tube of the FastDNA^®^ Spin Kit for Soil always recovered more DNA than the other bead preparations containing either 0.1 mm or 1.4 mm beads. Furthermore it was noted that although there was no statistically relevant difference between the use of 1.4 mm or 0.1 mm beads, the results changed, showing a statistical significance, when G2 was added (Table in S5 “d” Supporting Information).

**Fig 2.**
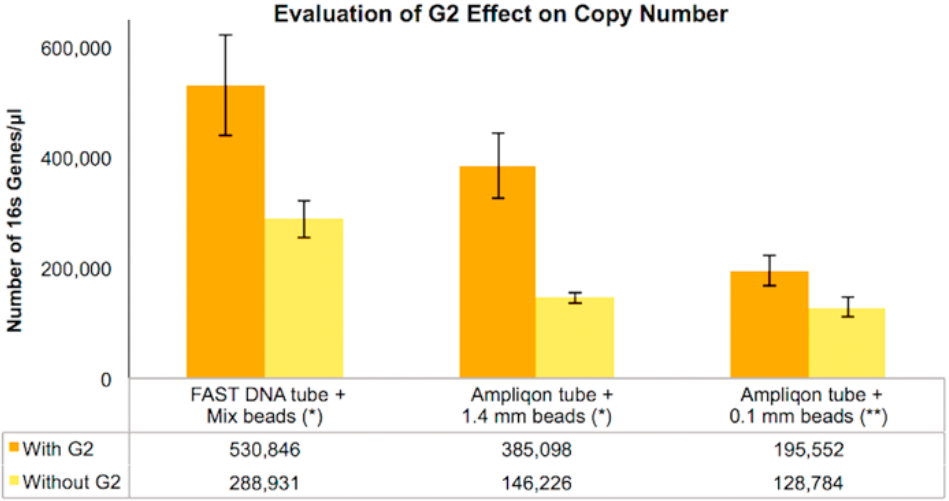
16S rRNA gene copy number variation after qPCR expressed as Genes/µl. A comparison between three commercial products with and without G2. Values shown are averages of n=3 technical replicates for n=3 biological replicates for each series. Error bars are calculated from the standard deviation for each series. (*) Significantly relevant comparison (Scheffe’s score > 26.117). (**) Not statistically relevant (Scheffe’s score < 26.117) (Table in S5 Supporting Information).

Furthermore, the results reported in Table in S5 “c” Supporting Information considered the possibility of basal contamination of the kit that may be relevant during the analyses or alternatively the possibility of a source of contamination introduced during the G2 freeze-drying process. In both cases, the control samples showed that basal contamination of the kit did not affect the results, and furthermore that the freeze-drying process did not introduce any more detectable DNA than the negative control.

To visualise this statistical evaluation based on all the different comparisons using Scheffe’s method, the following procedure was applied: Scheffe’s score (Table in S5 Supporting Information) was averaged by grouping different comparisons based on the variables examined, *i.e*. G2, bead size and tube plastic. The average score was then plotted and the results shown in Fig 3. Calculations are reported in Table in S6 Supporting Information. The results illustrated in Fig 3 show that the main influence to DNA yield came from the presence or absence of G2 in the lysing tubes, while the bead effect showed a smaller, but still relevant, significance. The results related to the two plastic tubes tested were non-significant.

**Fig 3.**
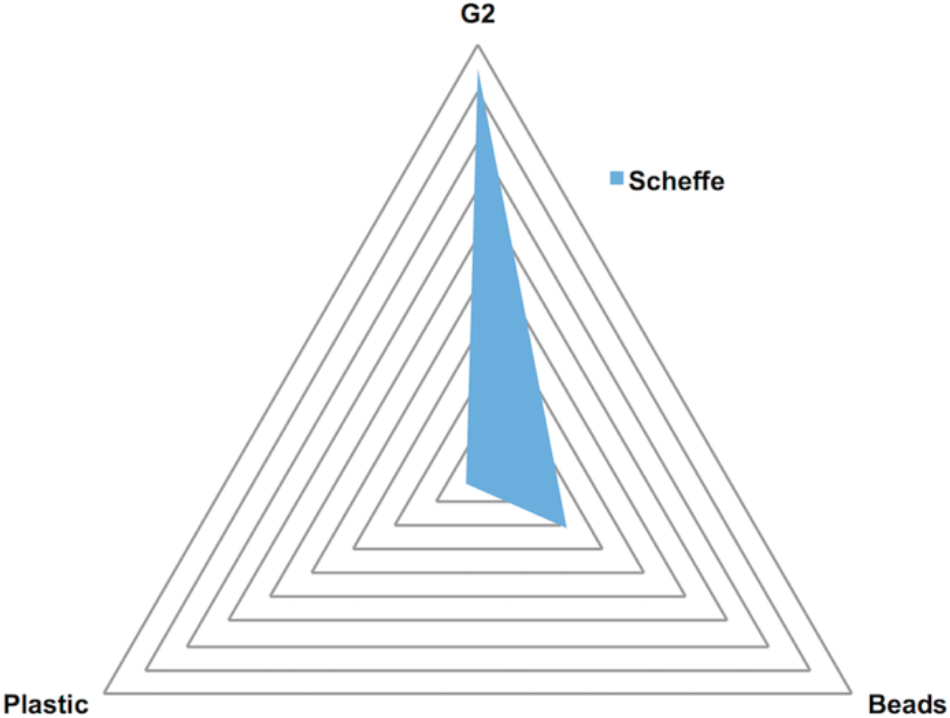
Visualisation of the average score of the Scheffe’s test. Visualisation of the average score of the Scheffe’s test of categories a, b and e, according to Table in S5 Supporting Information, visualising the impact of G2 compared with the effect of plastics and beads. The values on which this visualisation is based are reported in Table in S6 Supporting Information.

Considering that G2 is a DNA-based product, this could potentially affect the Qubit measurements since the signal recorded using a fluorescent dye is emitted when it binds to dsDNA. In such a situation there might be an overestimation of soil DNA from the Qubit result. To examine whether this was the case, the copy number variation of a genetic marker, such as the 16S rRNA gene for bacteria, was tested *via* qPCR. The results of the qPCR (Fig 2) confirmed the trend observed with the DNA yield provided by the Qubit analysis. G2 always allowed the recovery of the largest number of genes, while mixed beads also improved gene recovery (Scheffe’s score > 26.117 – Table in S5 Supporting Information).

The ANOVA based on the qPCR results described the dataset with a p-value of 1.0147*E^-^36 (Table in S4 Supporting Information). This means that in all the data produced by qPCR, taking into account the variance within and between groups, there was at least one difference that was statistically significant. However, since an ANOVA test does not identify where this difference is, Scheffe’s method was applied to compare the different combinations of G2, bead size and tube plastic (Table in S5 Supporting Information).

To visualise this statistical evaluation based on all the different comparisons using Scheffe’s method to show the impact of the single variable, the following procedure was adopted: Scheffe’s score was averaged (Table in S5 Supporting Information), grouping different comparisons based on the examined variables of G2, bead size and plastic tubes. The average score and the results are shown in Fig 3. Calculations are summarised in Table in S6 Supporting Information. Fig 3 shows that the most influential parameter of all the variables considered was the use of G2.

### Influence of G2 on the bacterial community structure

We also tested whether the use of G2 influenced microbial community composition, for example by enriching particular taxa or skewing the relative taxa abundance between samples. Hence, a subset of the DNA samples was sequenced for the V3-V4 region of the 16S rRNA gene. A statistical evaluation of the differences in alpha and beta diversity allowed an assessment of the impact of G2 on the microbial community DNA from this sample. Similarly, the impact of other variables, such as tubes and beads, on microbial community DNA composition was also evaluated. After data pre-processing through the QIIME2 pipeline, denoised and rarefied exact sequence variants were obtained. From these, the effects of G2 and bead size on alpha diversity were examined, focusing on two parameters: richness and evenness. These results are summarised in Fig 4.

**Fig 4.**
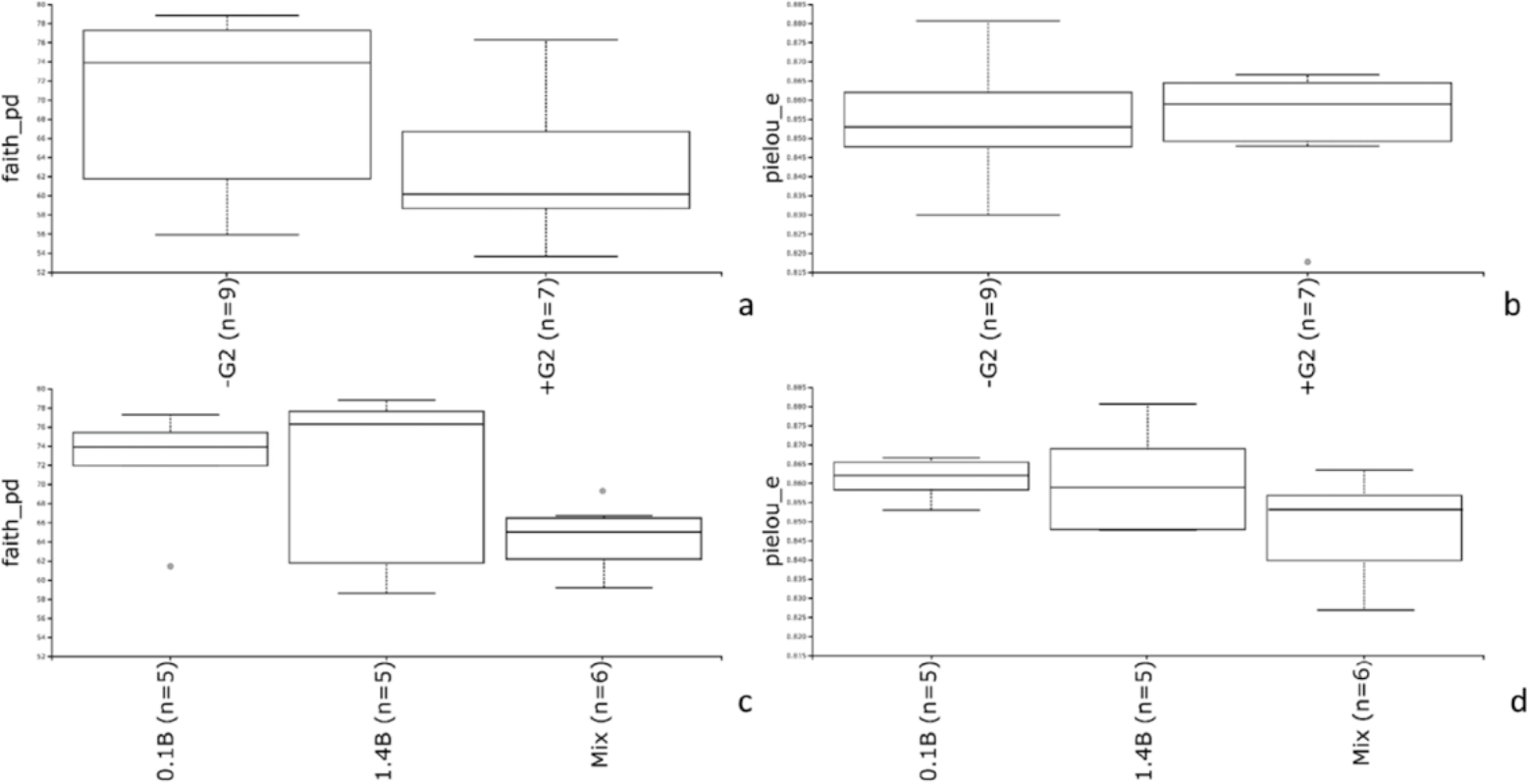
Alpha diversity analyses on G2 and bead impact on richness and evenness. (a) Boxplot based on the Faith-pd index and comparing all the samples with G2 (n=7) and without G2 (n=9). (b)Boxplot based on the Pielou-S score for evenness between samples obtained with G2 (n=7) and without G2 (n=9). (c) Boxplot based on the Faith-pd index and comparing all the samples with 0.1 mm (n=5), 1.4 mm (n=5) and mixed (n=6) beads. (d) Boxplot based on the Pielou-S score for evenness on samples obtained with 0.1 mm (n=5), 1.4 mm (n=5) and mixed (n=6) beads.

The presence of G2 did not statistically significant affect the richness (p = 0.08) or evenness (p = 0.874) of the microbial community, as shown in Figs 4A and 4B respectively. The same tests were applied to the results of the bead and tube comparison. While the use of two different plastic tubes did not show any statistically significant effect in terms of richness (p = 0.874) or evenness (p=0.322), this was not the case for the beads. In particular, the use of mixed bead sizes introduced a significant negative difference in richness when compared to 0.1 mm beads (p = 0.03), and in evenness when performing the same pairwise comparison with 0.1 mm beads (p=0.01). This change in the microbial community resulted in a smaller amount of sequence variants and evenness when mixed beads were used.

A further analysis was undertaken of the differences between the samples’ beta diversity using unweighted Unifrac and Bray-Curtis dissimilarity as metrics. These results were visualised using principal coordinates analysis (PCoA) plots and statistically evaluated in PERMANOVA. Fig 5 presents the PCoA plots showing the effects of the use of G2 (Fig 5A) and different beads (Fig 5B). These plots were obtained with the unweighted Unifrac distance matrix. The Bray-Curtis PCoA plot was consistent with this representation (not shown). PERMANOVA tests were performed with 999 permutations on beads, tubes and the G2 effect and the results are summarised in Table 2. These results showed that only mixed beads had a statistically relevant effect on beta diversity in the present study’s samples (p-value = 0.005 and 0.002). A PCoA plot of the tube type used did not allow a clear distribution to be distinguished between the two different tubes (not shown) and this was confirmed by the PERMANOVA analysis (p-value = 0.465) (Table 2). The PERMANOVA results are summarised in Table 2 with regard to the different factors evaluated and the comparisons performed.

**Fig 5.**
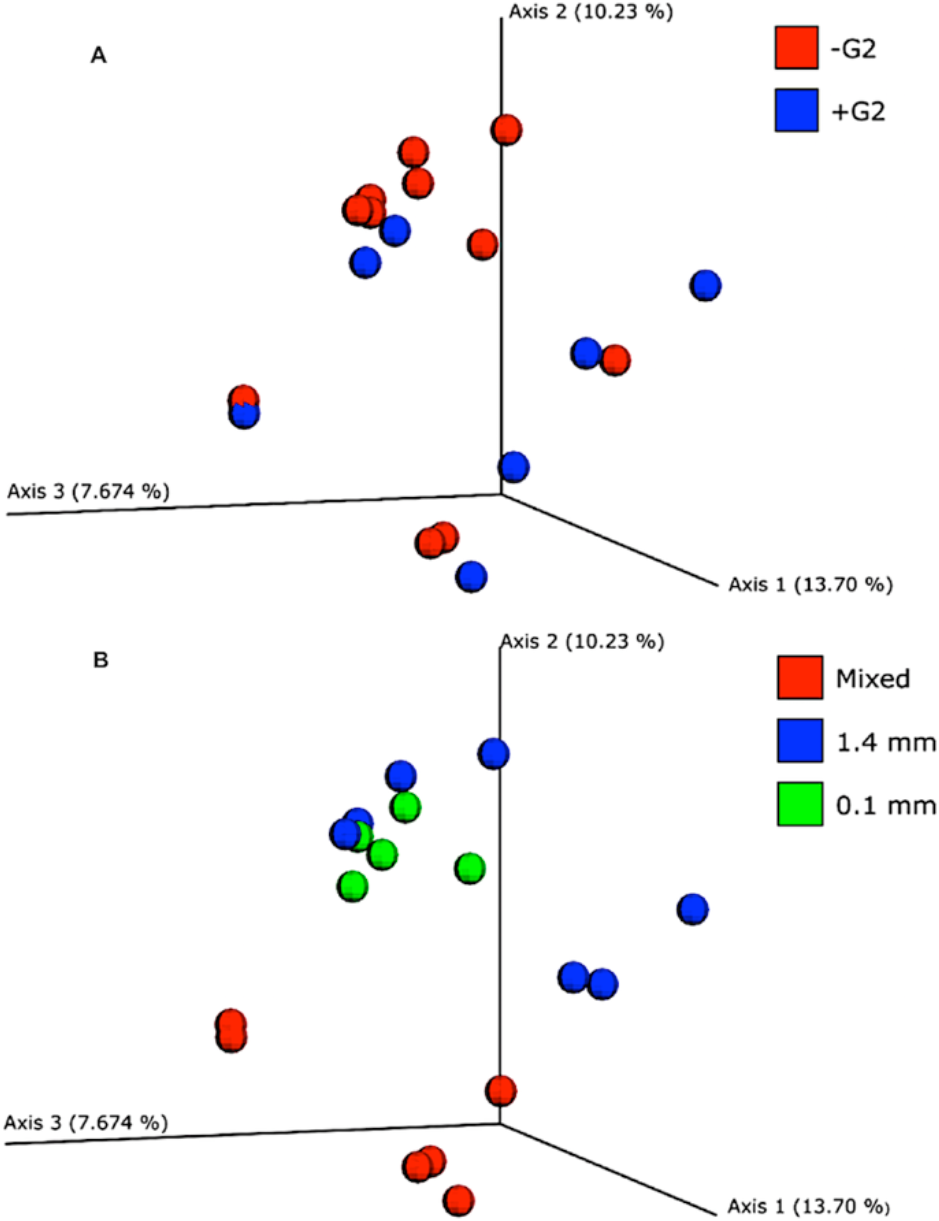
PCoA plots obtained from the UniFrac distance matrix of the sequenced samples from the different tube preparations. (A) Comparison of data obtained from samples with G2 (blue dots) and without G2 (red dots). (B) Comparison of data obtained from samples using 0.1 mm beads (green dots), 1.4 mm beads (blue dots) and mixed beads (red dots).

**Table 2.** Results of the PERMANOVA analyses.

| Variable | Groups | Pseudo f- | p-value | q-value |
| --- | --- | --- | --- | --- |
| G2 | Present /Absent | 0.979 | 0.499 | 0.499 |
| Beads | 0.1 mm / 1.4 mm | 0.907 | 0.787 | 0.787 |
| Beads | 0.1 mm / Mixed | 1.639 | 0.005 | 0.0075* |
| Beads | 1.4 mm / Mixed | 1.715 | 0.002 | 0.006* |
| Tube | Ampliqon/MP-Bio | 0.979 | 0.465 | 0.465 |

The first row refers to the use of G2, the second, third and fourth rows refer to the comparison of different groups of beads, and the fifth row refers to the tube effect. \*Statistically relevant comparisons. Pseudo f- refers to the ratio between cluster variance and within cluster variance; p-value is considered statistically relevant below 0.05; q-value is an adjusted p-value for false discovery rate (FDR) when multiple comparisons are performed.

The compositional bar chart made on phylum resolution (Fig in S7 Supporting Information) shows that the bacterial communities of the DNA extracts with and without G2 were not visually distinguishable. By contrast, when looking at the profiles using different types of beads, these differences were more evident, especially between the mixed beads compared with 1.4 mm or 0.1 mm beads. All the data reported so far were obtained by excluding the negative control from the alpha and beta diversity analyses in order to maximise the effect of the different variables reducing any false negative results. This could be done after checking the composition of the negative control through a taxa-bar plot. The dominant taxa in the negative control belonged to the genus *Ralstonia*, already reported and known to be common contaminants of kits and PCR reagents [45]. This taxon was filtered out of the other sample since it was present in low amounts (~0.1 % across the different series). Furthermore, the other taxa that appeared in the negative controls were checked and since they were dominants in all the samples and represented only a small fraction of the negative control, it was decided not to remove them. The taxa composition (Fig in S7 Supporting Information) of the microbial community in the soil horizon between 1.00 and 2.20 mbgl was shown to be mainly composed of members of the phyla of *Proteobacteria, Actinobacteria*, *Acidobacteria* and *Chloroflexi*. *Archaea* were also present and the dominant phylum was *Crenarcheota*, representing 10 % of the total relative abundance. For a better visualisation of the differences in terms of phyla attributable to the different variables involved, Fig 6 shows a heatmap of the microbial community composition of the dataset. Looking at the dendrogram on the left of Fig 6, two main groups can be distinguished according to the beads in the second heatmap. The use of mixed beads in particular altered the community profile, whereas none of the other variables produced this effect.

**Fig 6.**
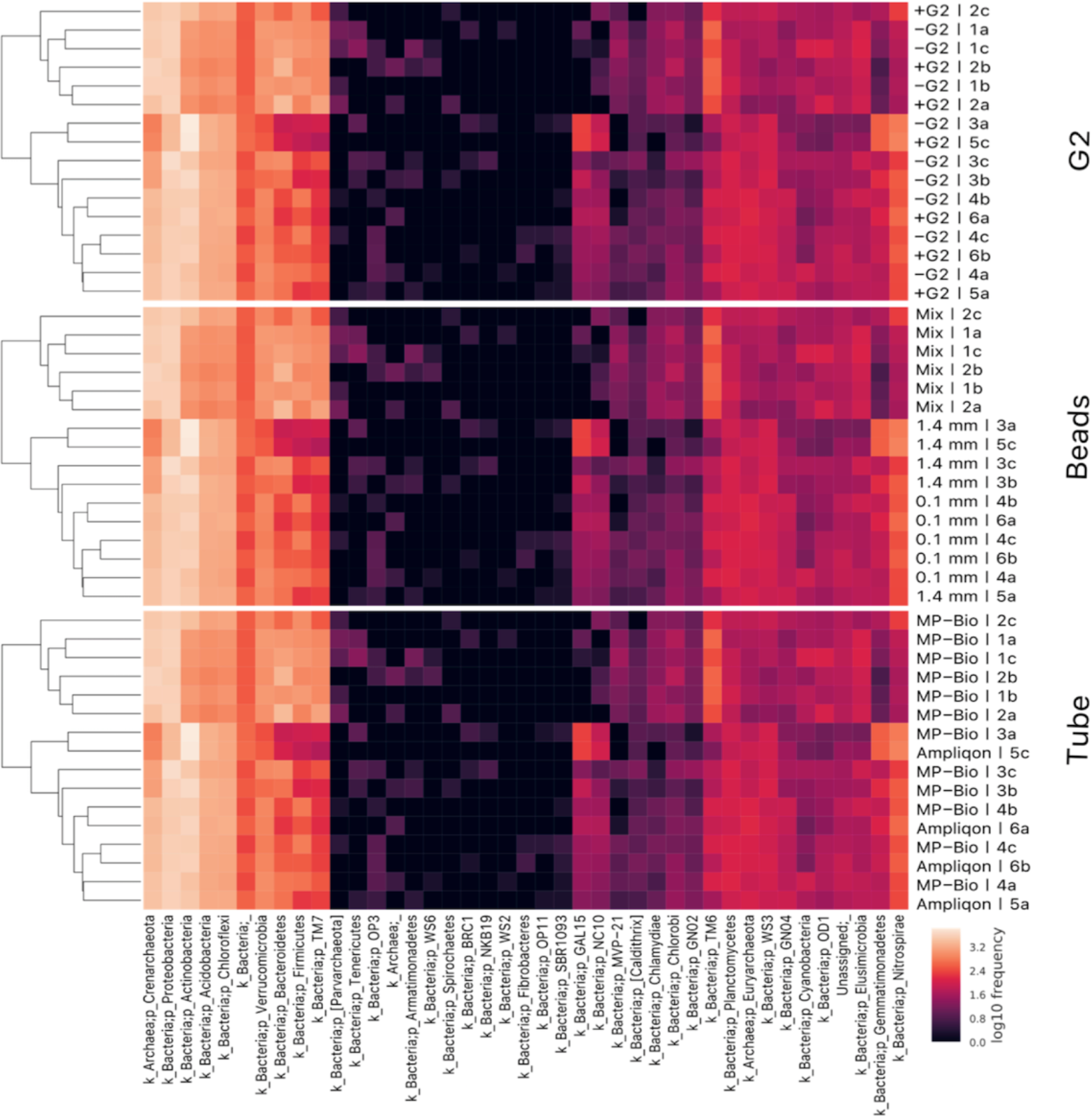
Heatmap at phylum level of the sequencing dataset. The logarithmic scale in which colour intensity determines the abundance of the taxa can be seen in the bottom right-hand corner. The top heatmap represents the samples grouped by presence/absence of G2, the middle one refers to the different bead sizes used, while the bottom one is related to the two different tube suppliers. The names of the samples are given on the right, while the names of the phylum are stated below. On the left the dendrogram of similarity between all the samples is presented.

Finally, an ANCOM test was run in order to identify differentially abundant microbes in the different groups of samples and determine which variables mostly affected the result.

Specifically, two different group comparisons were performed: one based on the presence/absence of G2 when mixed beads were used, and the other based on the difference between the use of mixed beads compared to 0.1 mm beads when G2 was used. These two tests confirmed that G2 did not statistically alter any taxa among the samples, as reported in Tables S8 and S9 Supporting Information. However there were four statistically relevant differences between the use of mixed and 0.1 mm beads. None of these four taxa belonged to the dominant fraction of the microbial community, but they represented around 1 % of the microbial community. Three of them were barely represented in the samples when 0.1 mm beads were used, while they were not present at all when mixed beads were used. They belong to the phyla of *Acidobacteria* and *Gemmatimonadetes*. Only one of these microbes was represented, at up to 1.36 %, when mixed beads were used compared to the 0.1 mm beads and it belongs to the order of *Legionellales*. The results of this last test are summarised in Tables S8 and S9 Supporting Information.

## Discussion

The use of G2 allowed the recovery of more DNA in all settings and samples, making it the largest impact variable compared to the other variables of interest (tubes and beads). The increased DNA yield found in the presence of G2 was in accordance with Bælum, Scheutz (46) and Jacobsen, Nielsen (47). The recovered DNA in the first case [46] was several orders of magnitude higher, probably related to a higher clay content and/or the clay mineral type increasing the retention capacity of the matrix on the released DNA. A 7.5-fold average increase in DNA yield was obtained in the second study, which was an inter-laboratory test [47].

To resolve whether the amount of DNA detected was derived from the microorganisms dwelling in the soil and not to any residual G2, the impact on 16S rRNA gene copy numbers was measured using qPCR (Fig 2). The output confirmed the same trend observed for the Qubit results, meaning a significantly higher recovery of bacterial genes in all tested cases. The combined use of qPCR and Qubit guarantees the best qualitative and quantitative analyses of DNA [46, 48]. The differences between the measured DNA amount and the gene copy number can be explained by the qPCR only being applied to the bacterial population, leading to an underestimation of the total number of cells in the samples. This could partially be compensated for by the copy number variations of the 16S rRNA gene in different bacterial populations possibly leading to an overestimation of the recovered amount of cells within the present samples.

G2 allowed the recovery of the largest amount of DNA when combined with mixed beads, but a relevant increase could also be observed by using just 1.4 mm beads, regardless of the plastic tube used. The number of cells could be estimated from the 16S rRNA gene numbers using the same methodology as Vishnivetskaya, Layton (24), in which it was assumed that the average 16S copy number for the microbial community was 3.6, based on the observation of Klappenbach, Saxman (49). This led to the calculation of a maximum of 3.7*10^5^ cells g^-^1 of soil using G2 and 2*10^5^ without G2. This value was lower than other reported cell counts, such as 2.2*10^8^ using the same kit, but in topsoil [24], a difference that can be attributed to a deeper sampled layer. Comparing the present results with a direct DNA extraction from a deep soil layer, such as the one performed by Taylor, Wilson (50), a more similar value is obtained of around 1.2*10^6^. Differences here could be related to the different soil type and the different kit used for the DNA extraction.

The combination of the two statistical tests, ANOVA and Scheffe, allowed multiple comparison analyses on a single dataset, as reported by Mermillod-Blondin, Rosenberg (51) and Brown (52). In addition to what is reported in Table in S5 Supporting Information and visualised in Fig 3, it was interesting to note that although there was not a statistically relevant difference between the use of 1.4 mm or 0.1 mm beads, the results changed when G2 was added and showed a statistical significance (Table in S5 “d” Supporting Information). This conversion might be due to the fact that 1.4 mm beads allowed improved lysis of the cells in the sample, probably because of a better dissolution of the soil particles, compared with 0.1 mm beads. However, if G2 was not added, most of the extracted DNA was suddenly adsorbed to the clay particles of the matrix, preventing its recovery.

To verify whether the use of G2 influenced the microbial composition, a subset of the DNA extracted from the soil samples was sequenced for the 16S rRNA gene. An evaluation of alpha diversity showed that evenness and richness did not change when comparing samples obtained with and without G2 or when using different tubes. Non-residual contamination of G2 has also been confirmed by Jacobsen, Nielsen (47), where in-depth sequencing did not show traces of the DNA originating from G2 in the final DNA extract [47]. In contrast, bead size type had a statistically relevant effect on these two parameters. The lower richness and evenness when mixed beads were used could be due to the fact that most of the total DNA extracted came from the dominant fraction of soil bacteria. Since these dominant bacteria were not selectively lysed by the 1.4 and 4.0 mm beads alone, but were by the 0.1 mm beads, this ratio between dominant and rare taxa would be maintained with every bead preparation, although the absolute number of lysed cells would be different. However, the library size from each sample was not proportional to the starting amount of DNA, leading to an uneven representation of rare taxa in the sample with a higher starting amount of DNA. Since the reads belonging to the dominant bacteria were sequenced multiple times for each sample, the richness when mixed beads were used was slightly lower. This was a common issue in all the PCR-based surveys and has also been confirmed by Gonzalez, Portillo (53). The lower evenness was also expected. In this case, samples from mixed bead tubes had a lower number of rare taxa, leading to a greater imbalance between dominant and rare taxa and thus to less evenness. With a larger amount of reads per sample, the effect on richness and evenness could probably be cancelled out or reverted due to the sequencing depth effect [54]. Some of the differences caused by uneven sequencing coverage could be reduced by performing a rarefaction on the sample, as was done in this study, but cannot be avoided completely [55].

In terms of beta diversity, only the use of mixed beads had an effect on the different samples. The PCoA distributions presented in Fig 5A show a clear cluster belonging to extracted mixed bead samples. The three bead sizes (0.1 mm, 1.4 mm and 4.0 mm) could act together to provide a better dissolution of the sample and recover more DNA, but if sequencing is not deep enough it could lead to the detection of a reduced number of taxa, as was the case in this study. Furthermore, since commercial products were used here, it is worth noting that the amount of beads differed between the MP-Biochemical tubes and the Ampliqon tubes. In particular MP-Bio tubes have a higher volume of beads, and this could explain some of the differences detected in terms of the amount of DNA and the quality of the taxa. The synergistic bead effect, both positive and negative, was not evaluated in this study. Furthermore, the PCoA plots presented in Fig 5, and then confirmed by the PERMANOVA analyses (Table 2), showed that there was no statistical difference in microbial composition between the samples with or without the use of G2 (Fig 5A) and also irrespective of the plastic tubes used (not shown). Finally, in terms of taxonomic composition of the microbial community (Fig in S7 Supporting Information), these results are consistent with Janssen (56) and He, Guo (57) with regard to the dominant phyla of *Proteobacteria*, *Actinobacteria* and *Acidobacteria*in soil at different horizon levels.

## Conclusions

Based on DNA yield quantification and gene copy-number detection after qPCR, the current study demonstrated that the use of the commercial product G2 DNA/RNA Enhancer^®^ (Ampliqon A/S, Odense, Denmark) increased the amount of DNA recovered from composite and homogenised silty clay soil samples. Furthermore, the use of G2 did not introduce any significant differences in the richness or evenness of the bacterial community obtained after amplicon library sequencing when compared with those samples sequenced without the addition of G2. The two plastic lysing tubes tested had no effect on either the yield or the composition of the microbiota.

In contrast, the use of different bead sizes had a significant effect. A higher DNA yield was obtained with the simultaneous presence of differently sized beads. Moreover, the use of mixed beads in this case led to a slightly lower richness and evenness in the taxa distribution, an effect that could be explained by the sequencing depth. In terms of future perspectives coming out of this study, it would be worth applying G2 to other kinds of samples in which DNA recovery could be affected by proteins or other compounds biasing downstream application.

These tests may provide useful information for the improvement of existing commercial products in DNA/RNA extraction kits, raising awareness about the optimal choice of additives.

## Acknowledgments

We acknowledge Jesper S. Pedersen (Ampliqon A/S) for preparing and providing laboratorial material and Patricia Benitez for fieldwork support.

## Supporting information

**S1 Supporting Information.**
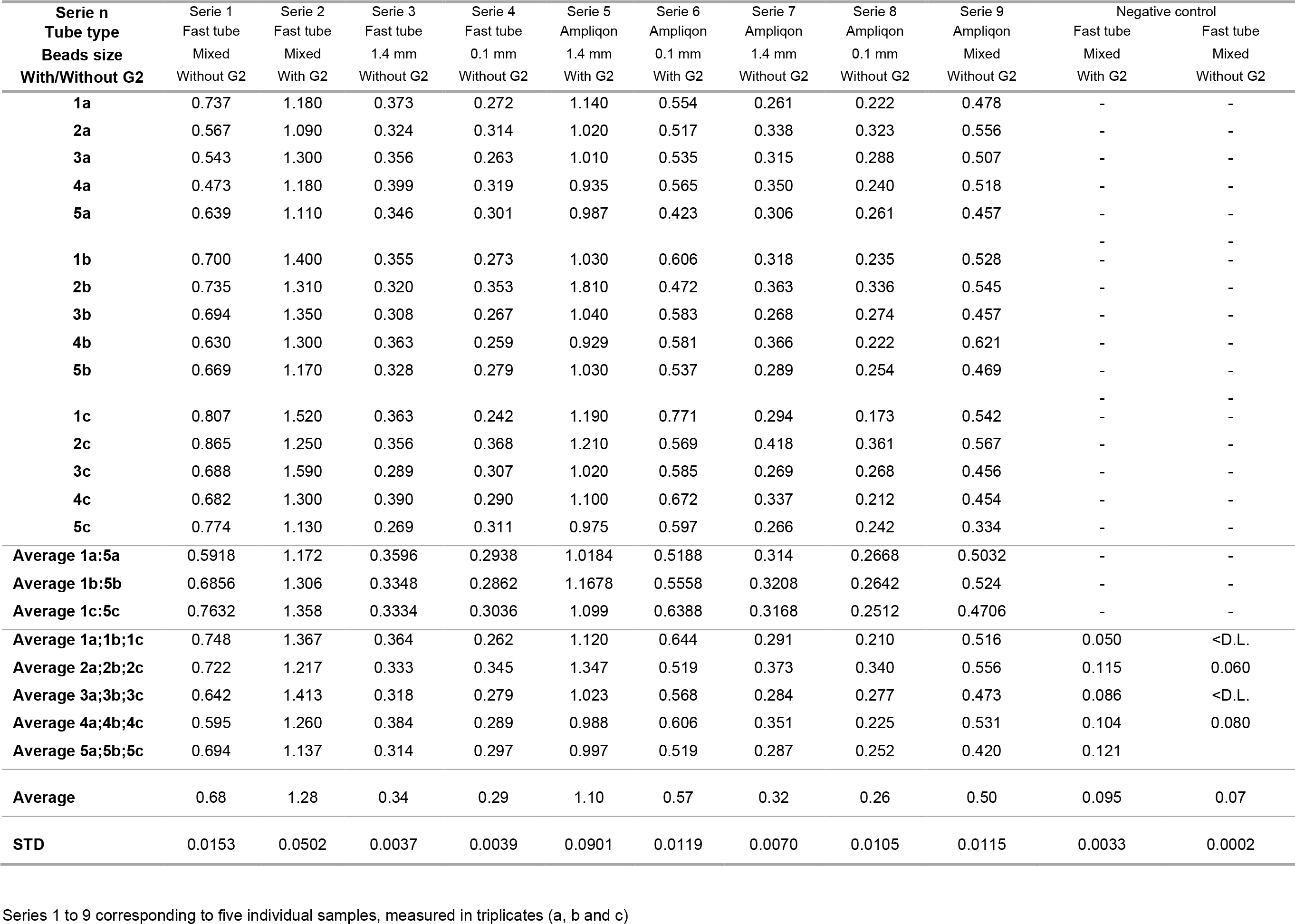
Qubit measurements (ng/µl).

**S2 Supporting Information.**
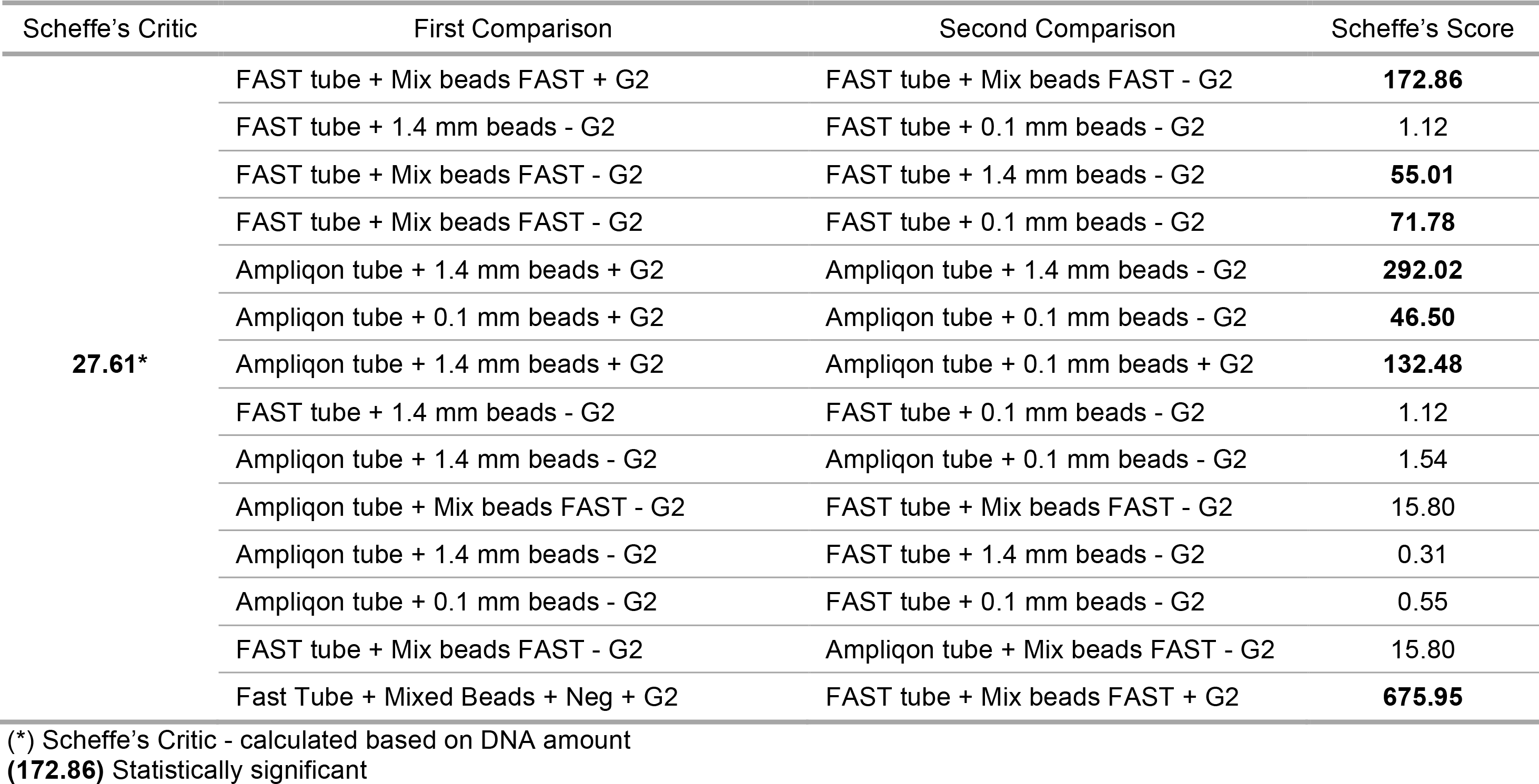
Scheffe’s score for Qubit dataset based on the difference series comparisons.

**S3 Supporting Information.** qPCR copy-number results (quantification based on 16s Gene/µl).

| FAST tube +<br>Mix beads<br>FAST - G2 | FAST tube +<br>Mix beads<br>FAST + G2 | FAST tube +<br>1.4 mm beads<br>- G2 | FAST tube +<br>0.1 mm beads<br>- G2 | Ampliqon tube<br>+ 1.4 mm<br>beads + G2 | Ampliqon tube<br>+ 0.1 mm<br>beads + G2 | Ampliqon tube<br>+ 1.4 mm<br>beads - G2 | Ampliqon tube<br>+ 0.1 mm<br>beads - G2 | Ampliqon tube +<br>Mix beads FAST<br>- G2 |
| --- | --- | --- | --- | --- | --- | --- | --- | --- |
| 239.674 | 533.510 | 150.081 | 136.344 | 344.158 | 197.511 | 146.488 | 122.612 | 306.625 |
| 288.782 | 474.536 | 146.095 | 119.145 | 361.691 | 217.290 | 147.103 | 99.747 | 278.049 |
| 278.633 | 373.218 | 156.656 | 86.502 | 447.643 | 139.696 | 135.779 | 104.346 | 257.049 |
| 357.835 | 441.625 | 120.722 | 168.695 | 460.500 | 174.806 | 168.359 | 161.059 | 289.533 |
| 310.236 | 711.104 | 161.245 | 126.655 | 417.669 | 188.429 | 134.722 | 141.191 | 286.099 |
| 324.357 | 541.500 | 98.555 | 114.850 | 436.516 | 190.285 | 142.007 | 143.750 | 271.976 |
| 265.700 | 547.510 | 154.716 | 126.569 | 383.342 | 246.825 | 139.759 | 128.784 | 266.401 |
| 271.138 | 608.738 | 158.943 | 78.312 | 347.810 | 193.563 | 143.349 | 128.784 | 275.977 |
| 264.020 | 545.876 | 169.024 | 89.805 | 266.555 | 211.566 | 157.162 | 128.784 | 278.869 |
| Average |  |  |  |  |  |  |  |  |
| 288.931 | 530.846 | 146.226 | 116.320 | 385.098 | 195.552 | 146.081 | 128.784 | 278.953 |
| Std. |  |  |  |  |  |  |  |  |
| 34.101 | 91.386 | 21.150 | 26.673 | 58.663 | 27.878 | 10.108 | 17.944 | 13.435 |

**S4 Supporting Information.**
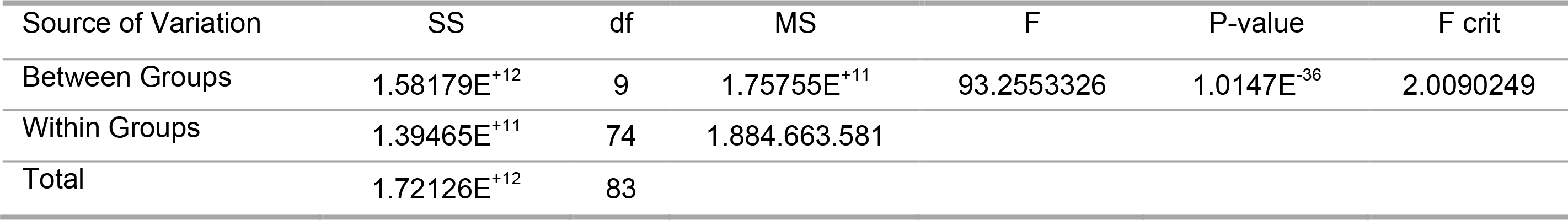
ANOVA table based on qPCR dataset.

**S5 Supporting Information.**
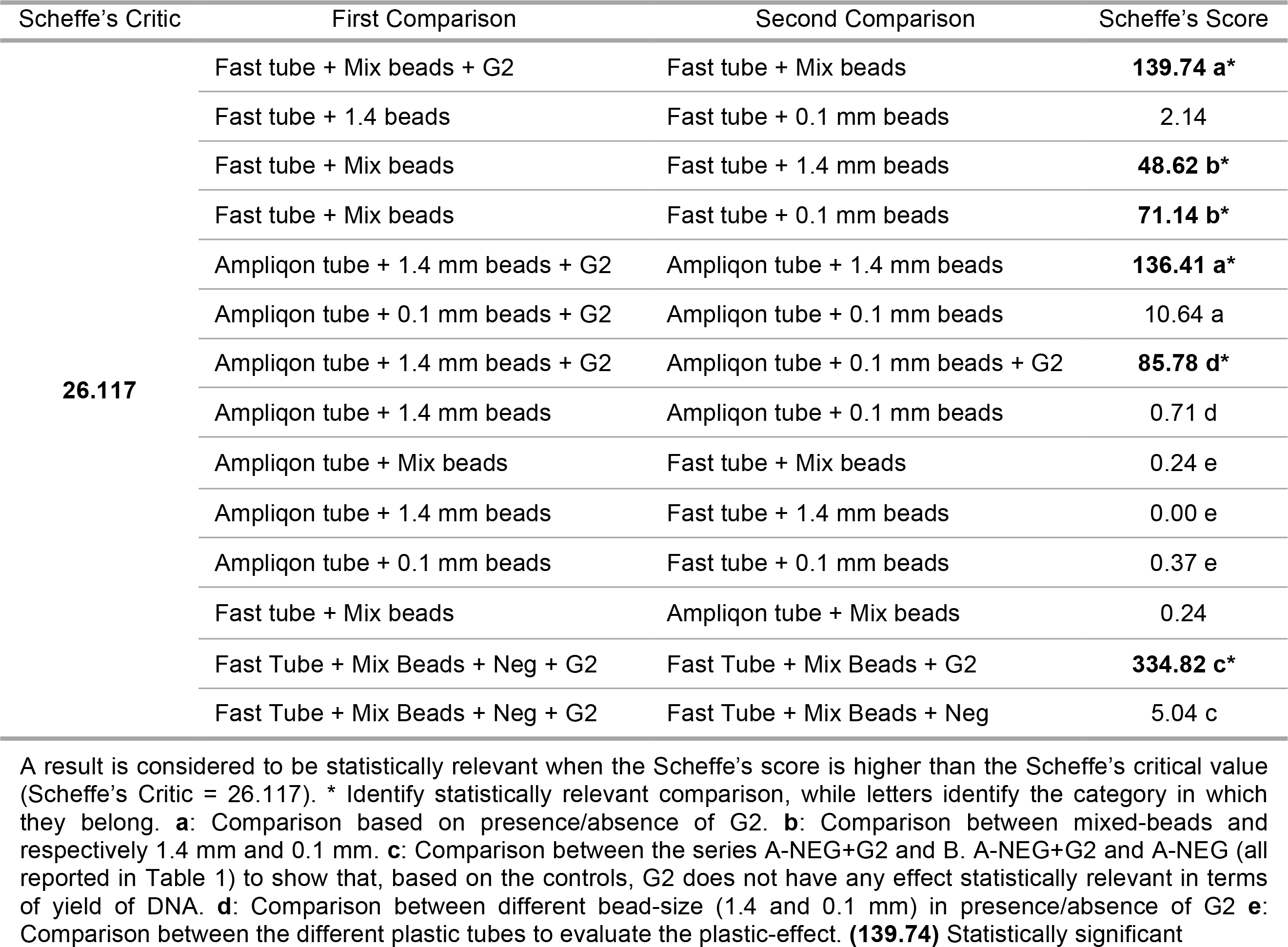
Scheffe’s score for qPCR dataset based on the difference series comparisons.

**S6 Supporting Information.** Scheffe’s test results as presented in Table S5 Supporting Information.

| **G2** | **Beads** | **Tubes** |
| --- | --- | --- |
| 139.74 | 2.10 | 0.00005 |
| 136.40 | 48.20 | 0.37 |
| 10.60 | 71.10 | 0.23 |
| **Averages** |  |  |
| 95.58 | 40.47 | 0.20 |

Data was used for Figure 3 preparation

**S7 Supporting Information.**
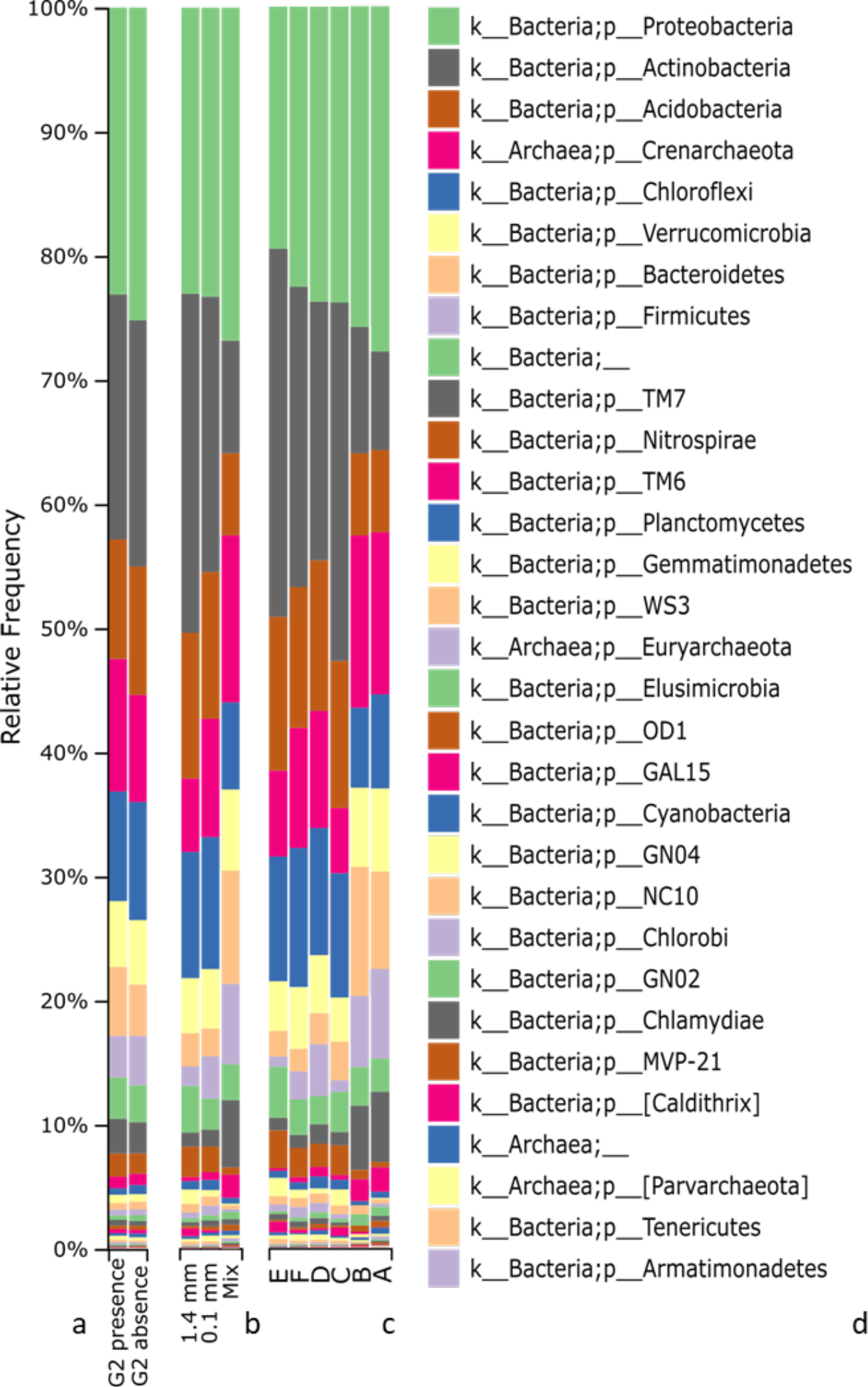
Compositional bar charts on phylum resolution. (a) G2 presence/absence, (b) beads-effect, (c) differences among Series, and (d) relative taxonomic legend. The chromatic order in the label starts from the top-most abundant and re-cycle every eight taxa.

**S8 Supporting Information.**
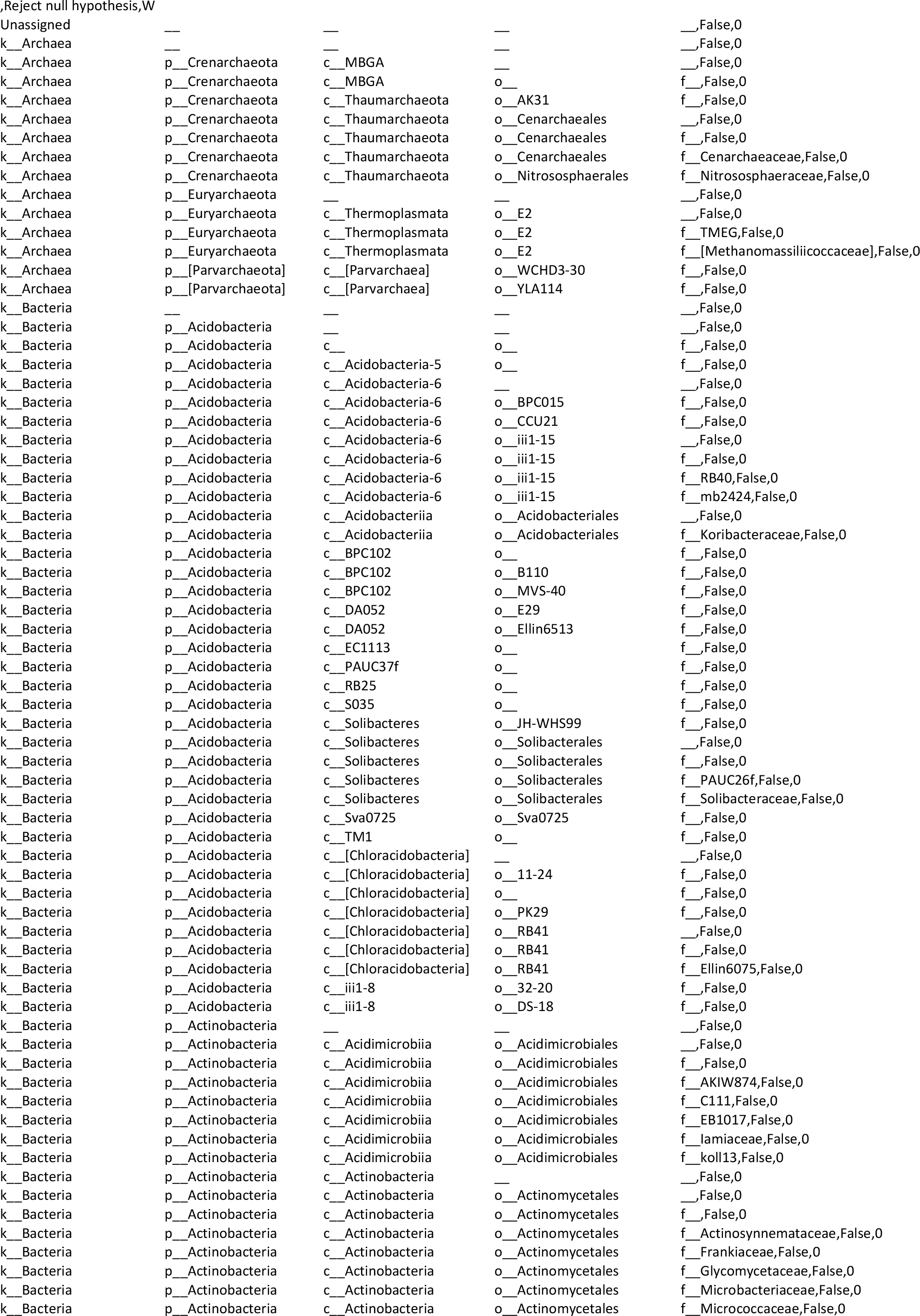
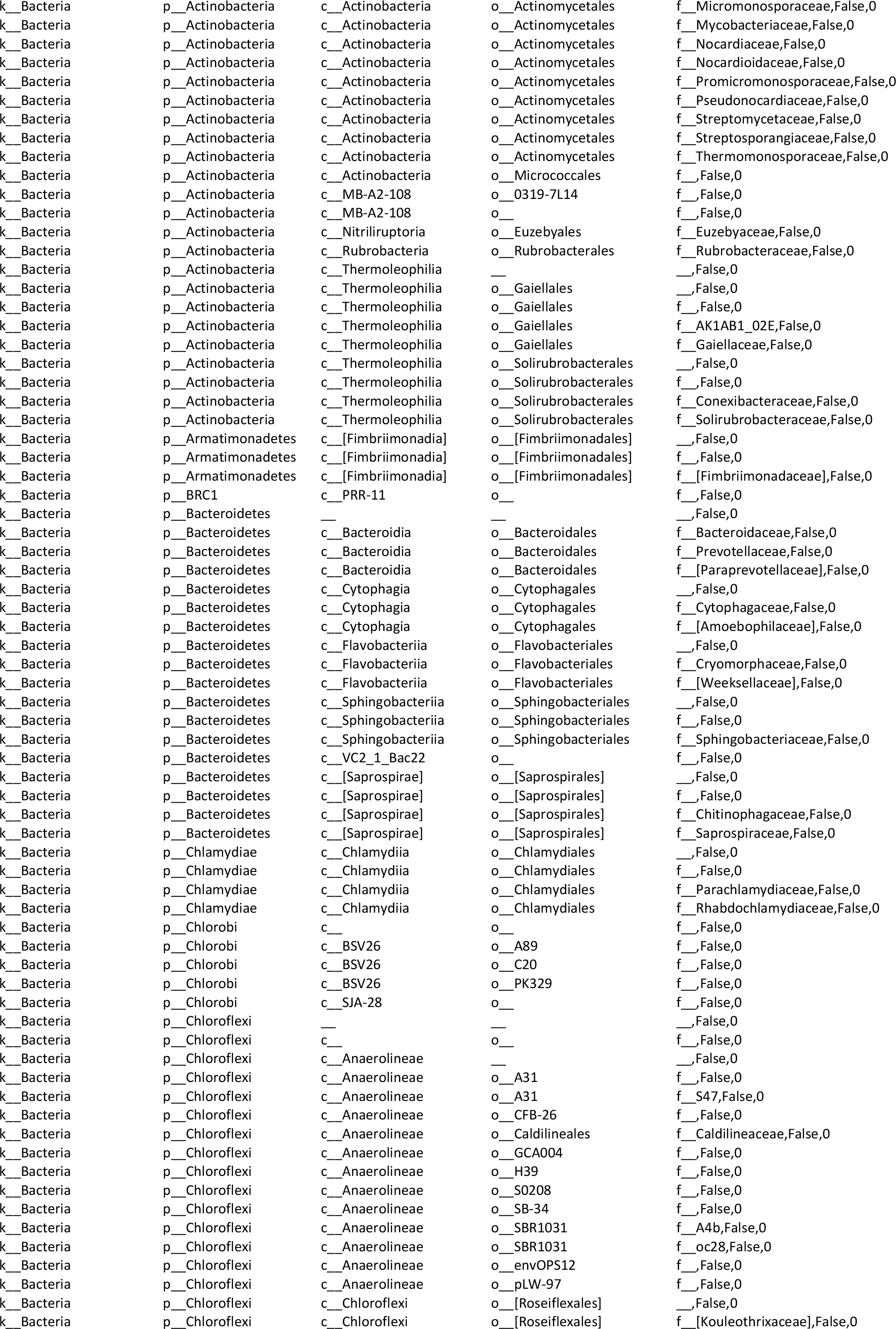
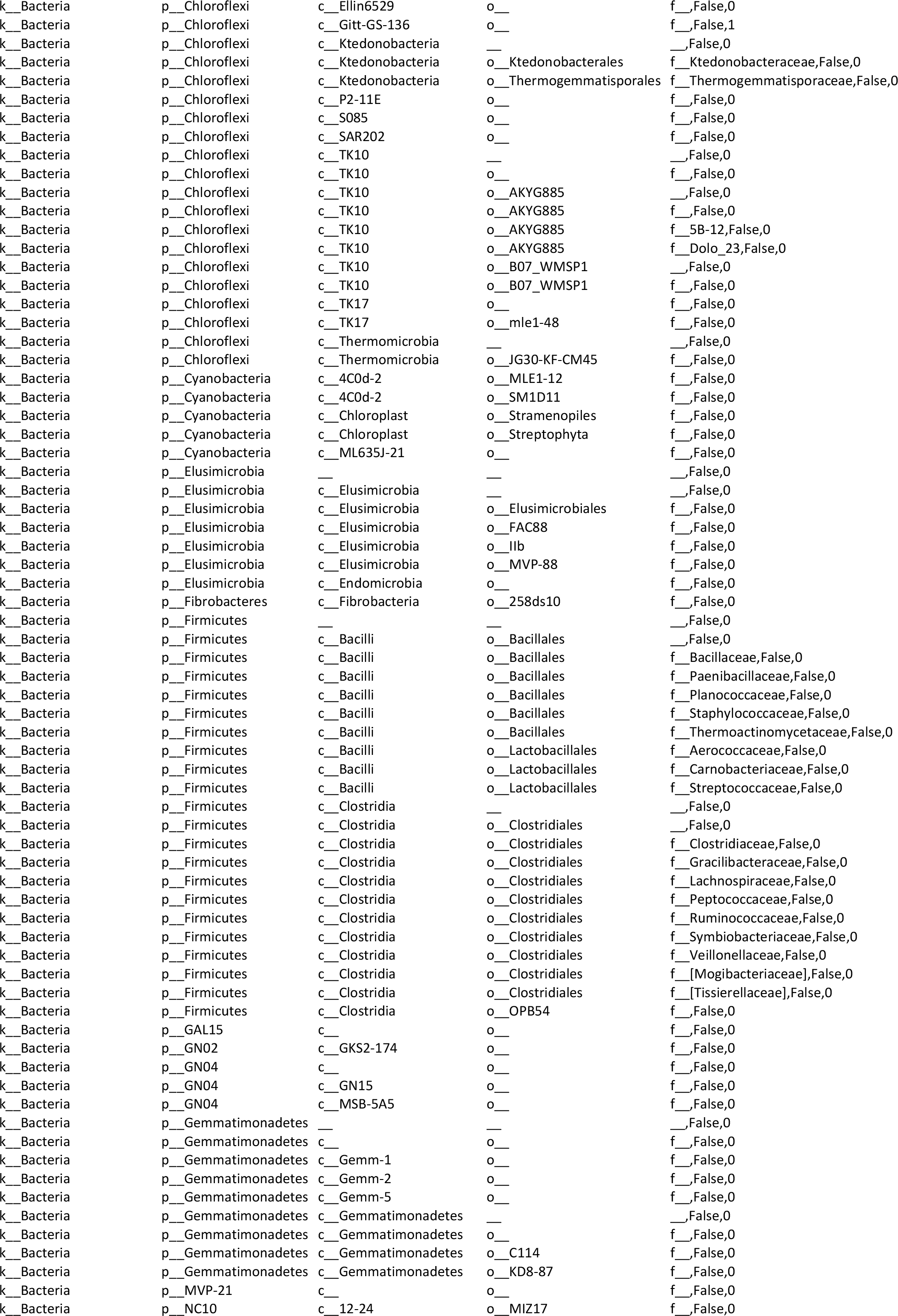
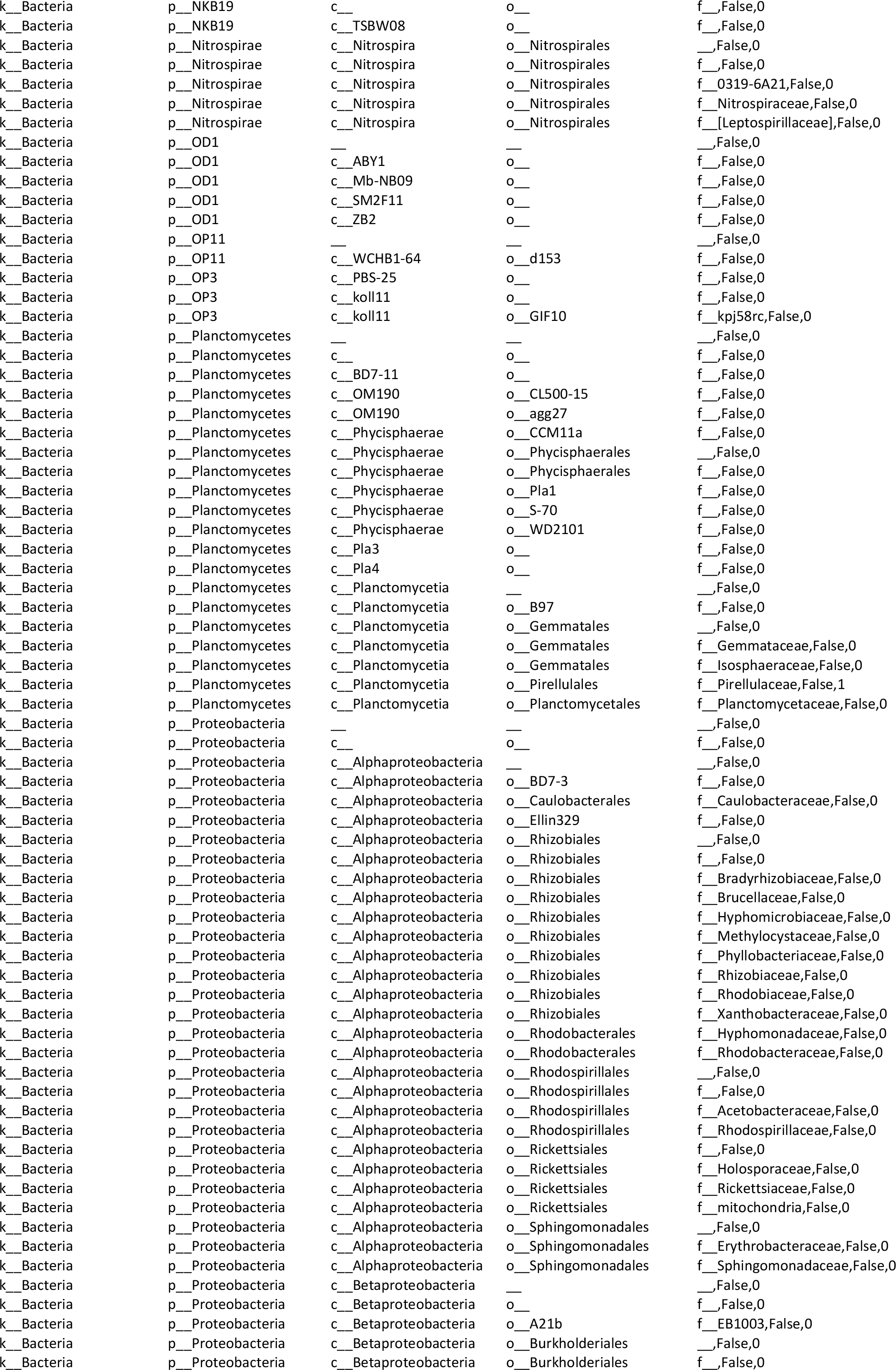
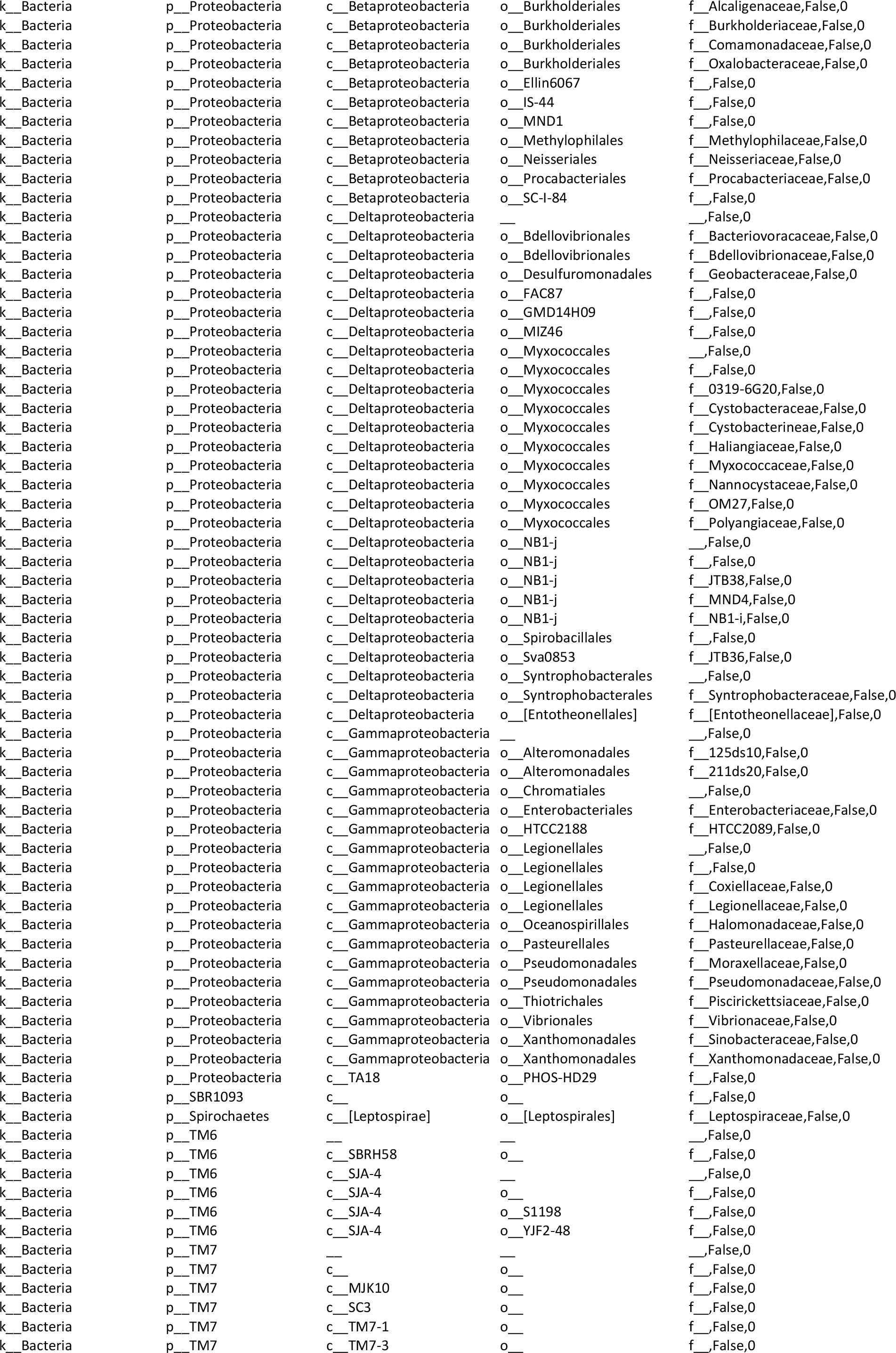
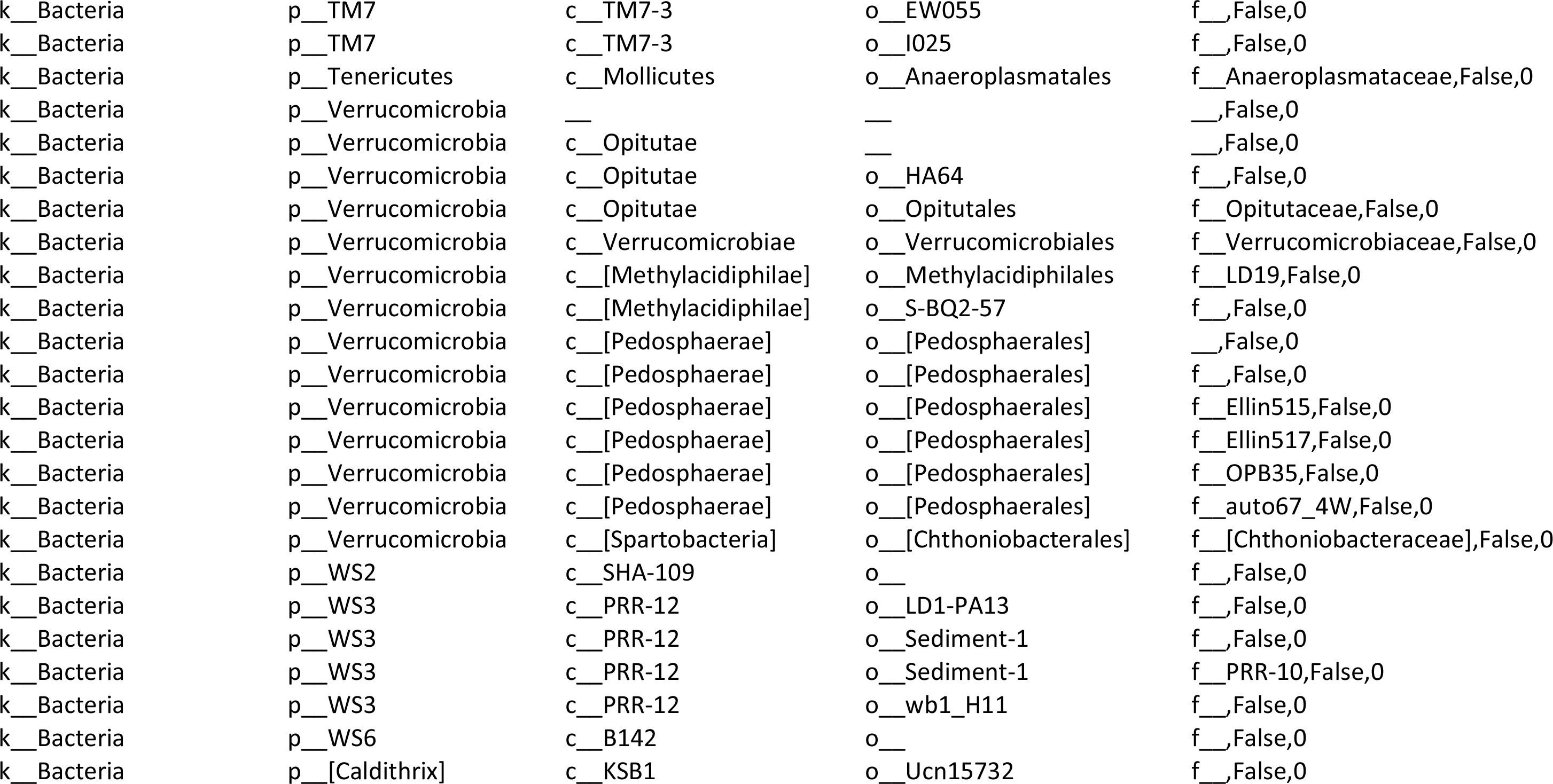
ANCOM summary based on G2 comparison.

**S9 Supporting Information.**
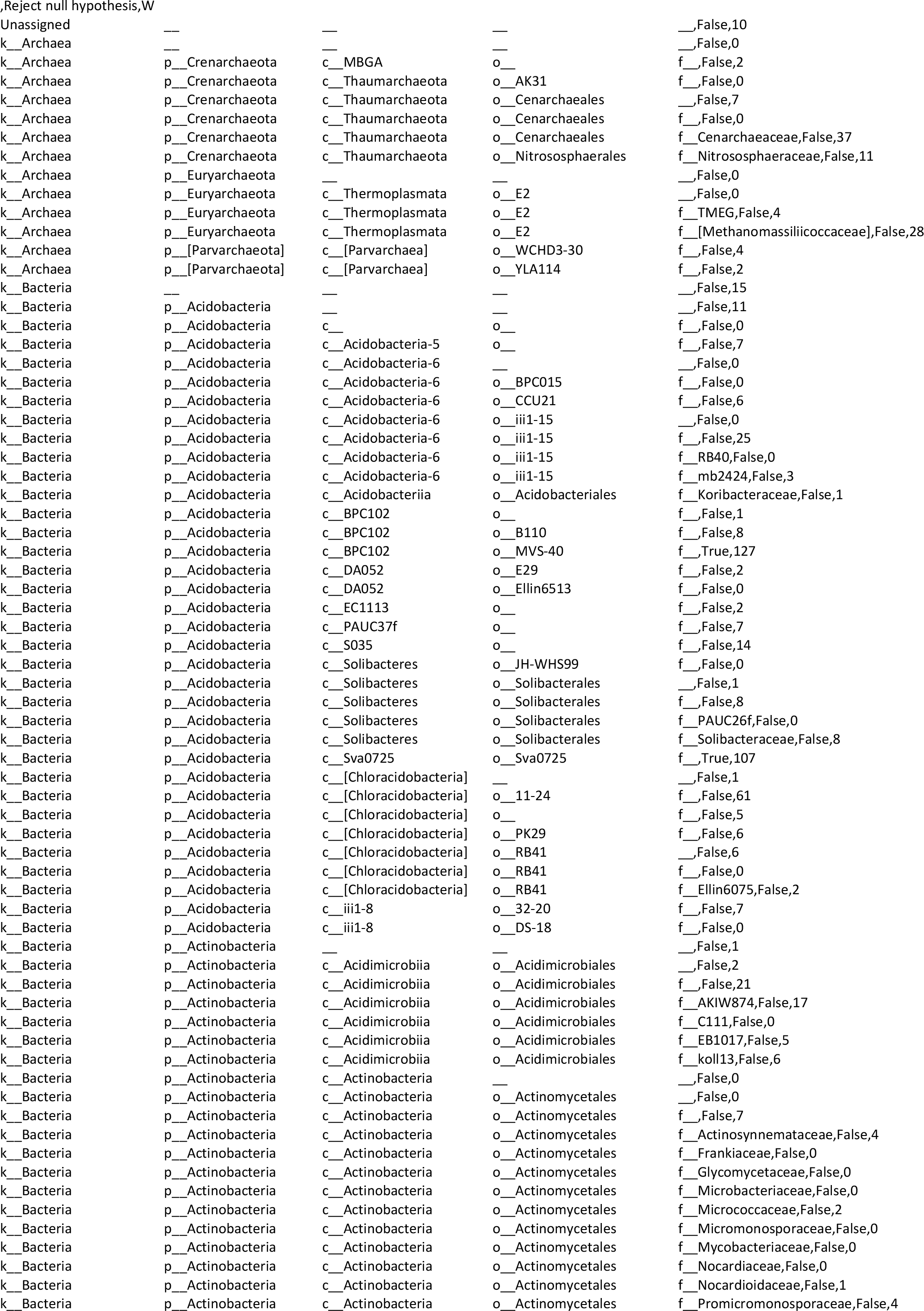
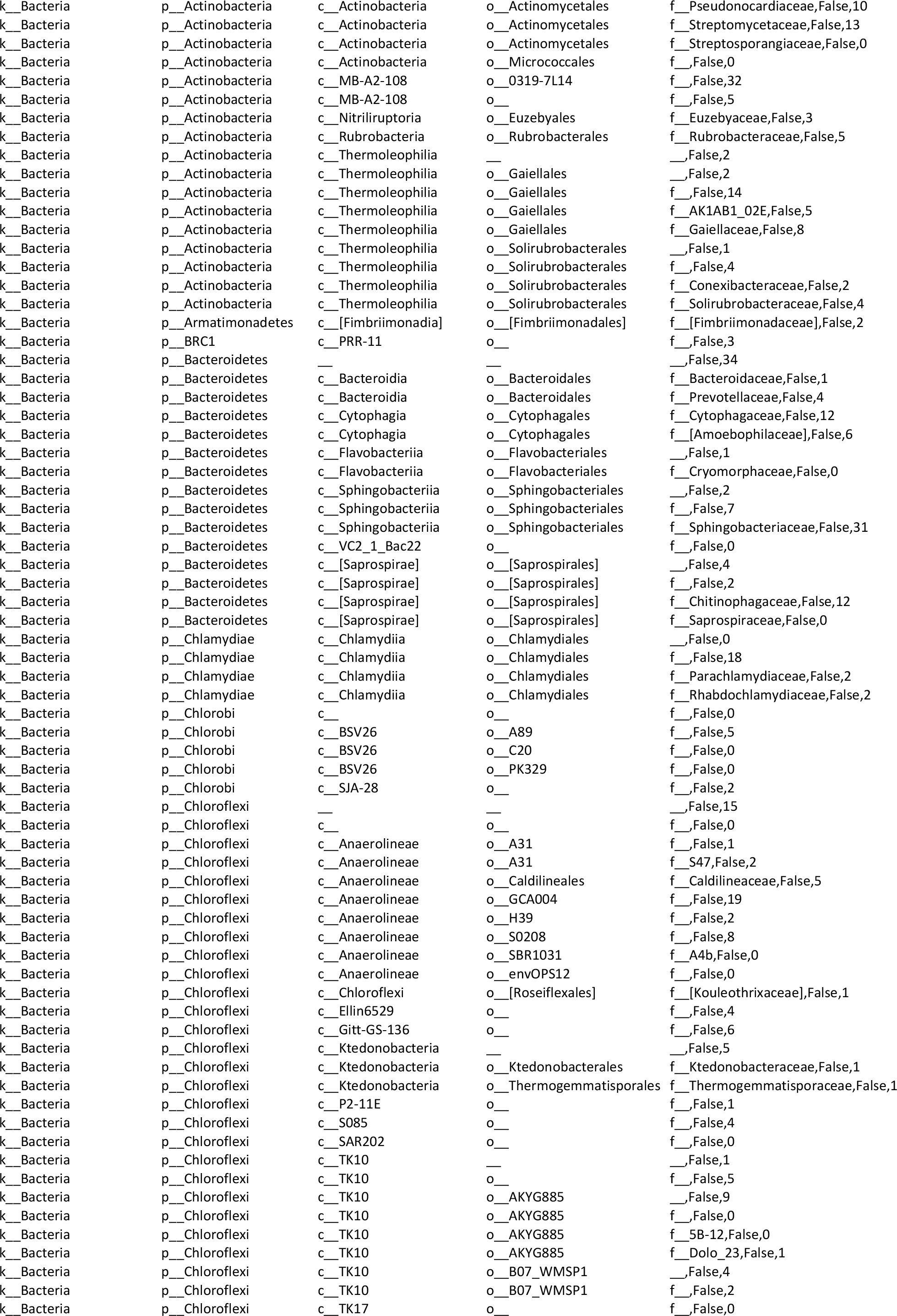
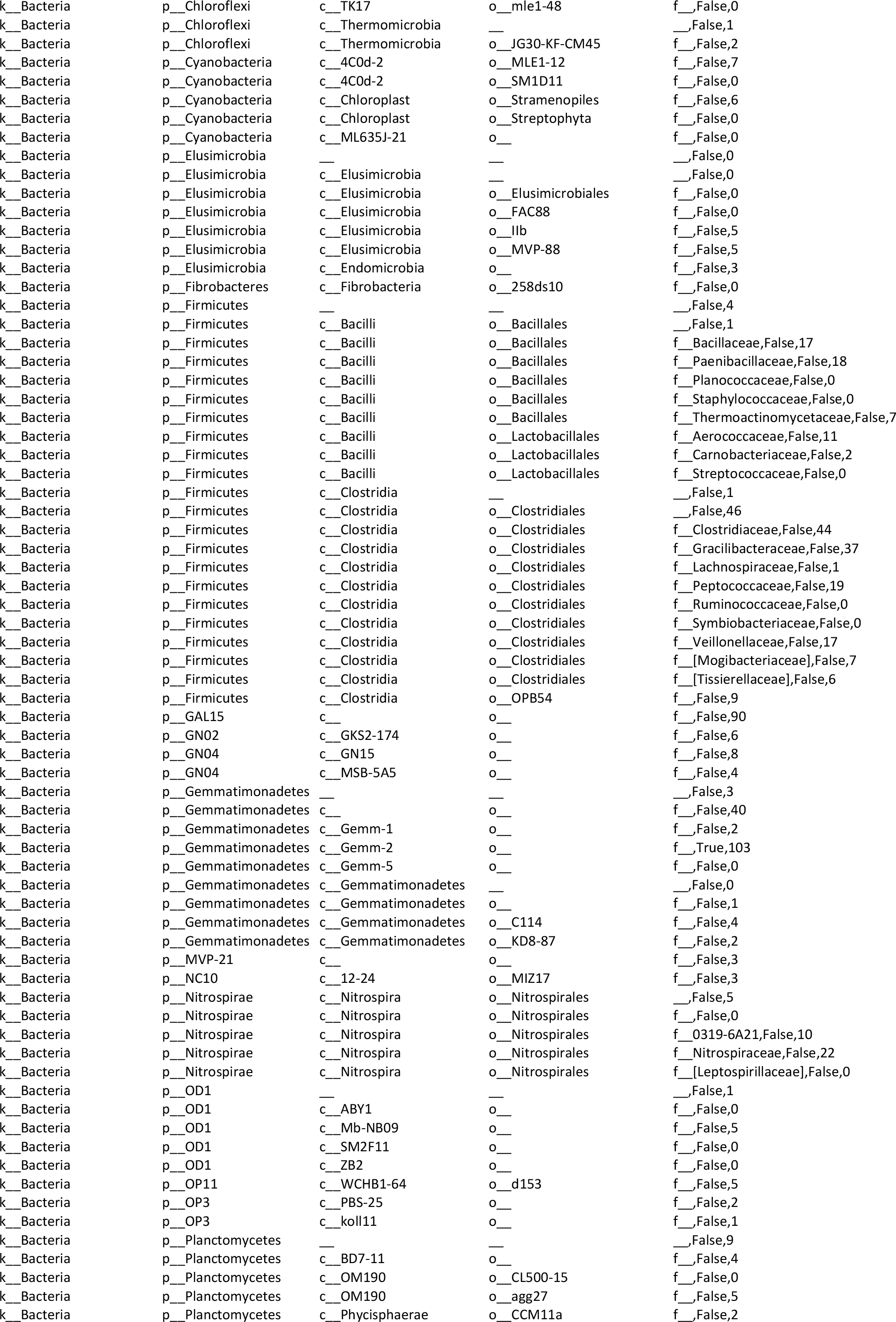
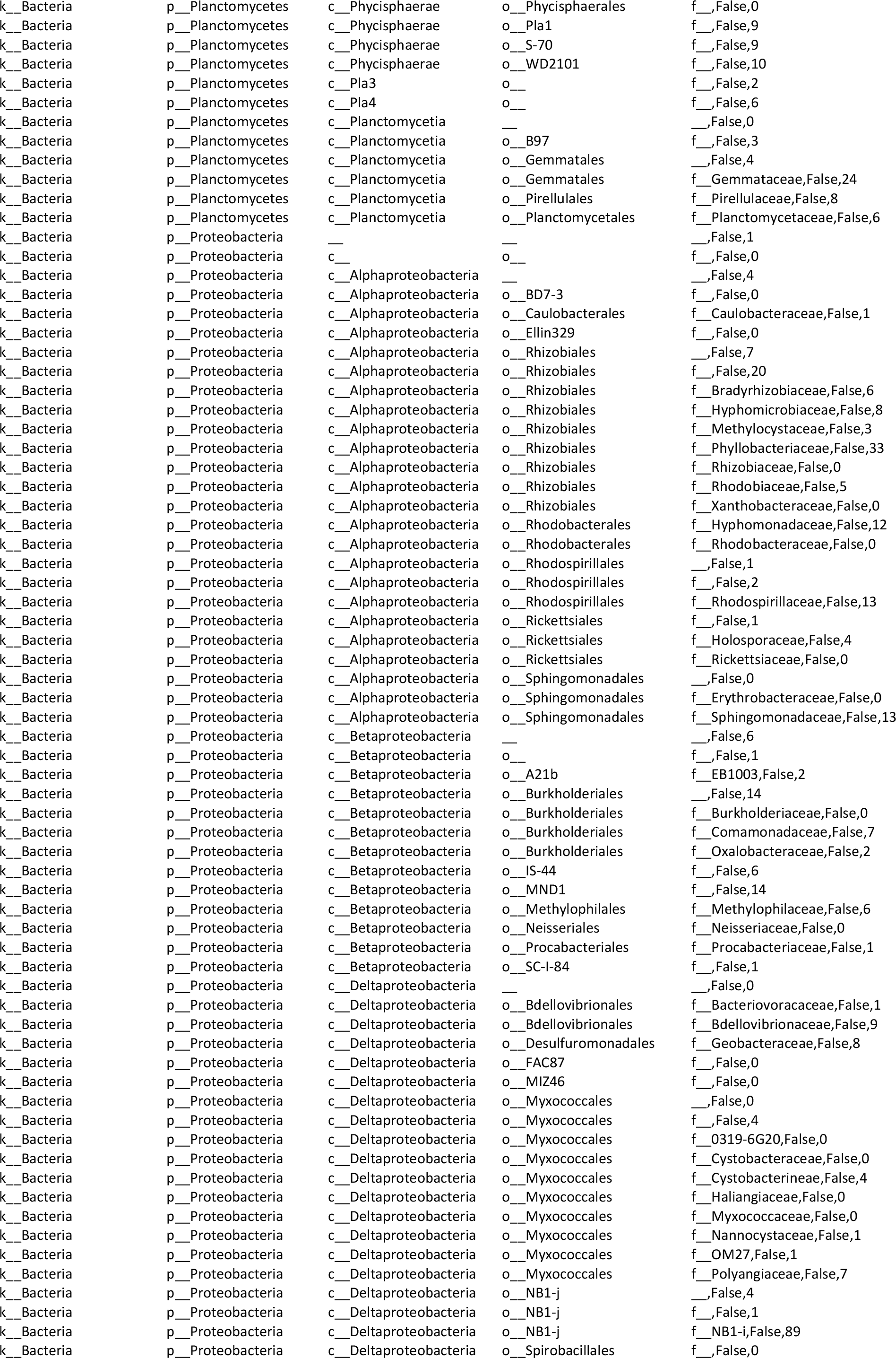
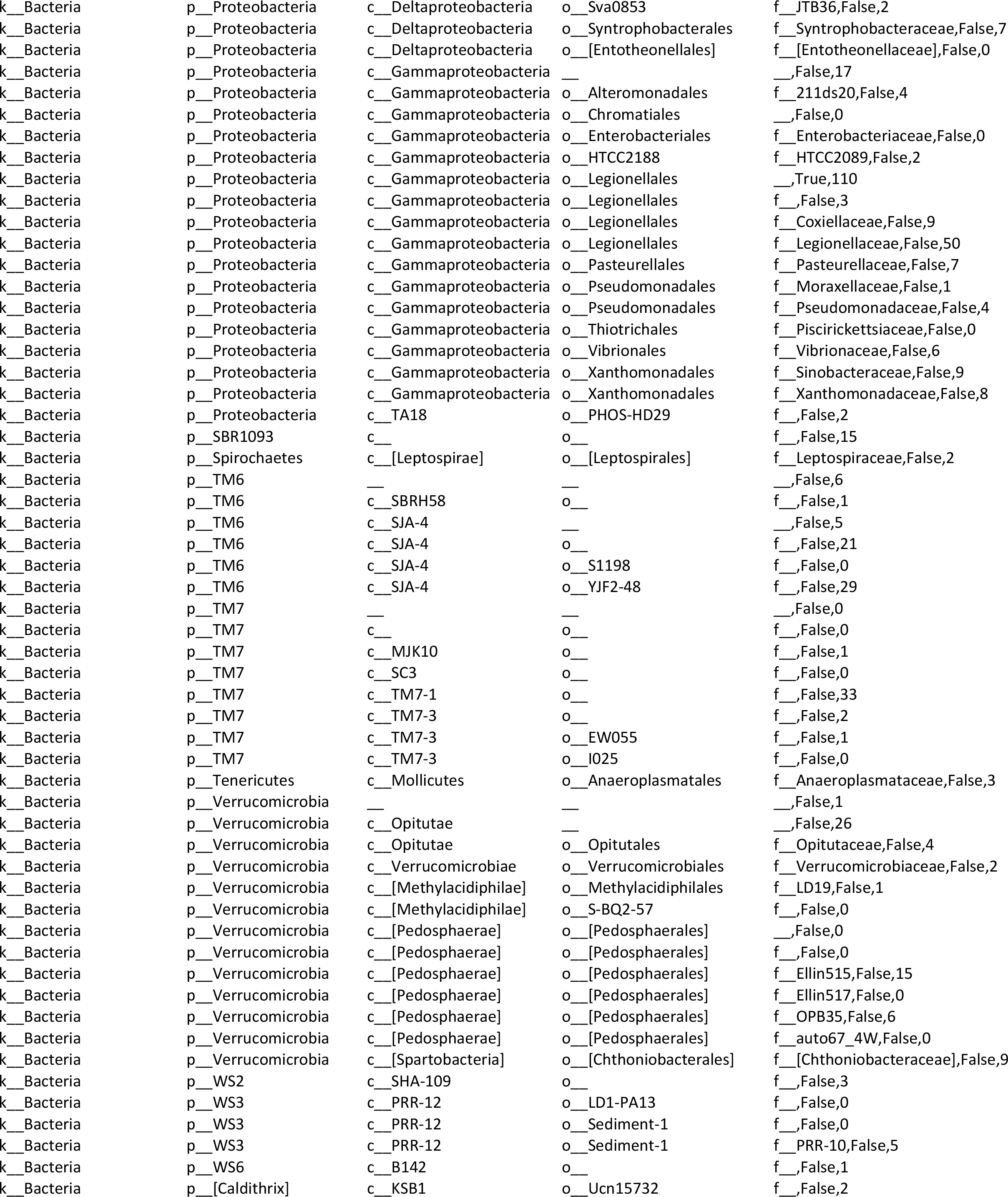
ANCOM summary based on beads comparison.

